# Modeling binary and graded cone cell fate patterning in the mouse retina

**DOI:** 10.1101/650606

**Authors:** Kiara C. Eldred, Cameron Avelis, Robert J. Johnston, Elijah Roberts

## Abstract

Nervous systems are incredibly diverse, with myriad neuronal subtypes defined by gene expression. How binary and graded fate characteristics are patterned across tissues is poorly understood. Expression of opsin photopigments in the cone photoreceptors of the mouse retina provides an excellent model to address this question. Individual cones express S-opsin only, M-opsin, or both S-opsin and M-opsin. These cell populations are patterned along the dorsal-ventral axis, with greater M-opsin expression in the dorsal region and greater S-opsin expression in the ventral region. Thyroid hormone signaling plays a critical role in activating M-opsin and repressing S-opsin. Here, we developed an image analysis approach to identify individual cone cells and evaluate their opsin expression from immunofluorescence imaging tiles spanning roughly 6 mm along the D-V axis of the mouse retina. From analyzing the opsin expression of ∼250,000 cells, we found that cones make a binary decision between S-opsin only and co-expression competent fates. Co-expression competent cells express graded levels of S- and M-opsins, depending nonlinearly on their position in the dorsal-ventral axis. M- and S-opsin expression display differential, inverse patterns. Using these single-cell data we developed a quantitative, stochastic model of cone cell decisions in the retinal tissue based on thyroid hormone signaling activity. The model recovers the probability distribution for cone fate patterning in the mouse retina and describes a minimal set of interactions that are necessary to reproduce the observed cell fates. Our study provides a paradigm describing how differential responses to regulatory inputs generate complex patterns of binary and graded cell fates.

**Author Summary:** The development of a cell in a mammalian tissue is governed by a complex regulatory network that responds to many input signals to give the cell a distinct identity, a process referred to as cell-fate specification. Some of these cell fates have binary on-or-off gene expression patterns, while others have graded gene expression that changes across the tissue. Differentiation of the photoreceptor cells that sense light in the mouse retina provides a good example of this process. Here, we explore how complex patterns of cell fates are specified in the mouse retina by building a computational model based on analysis of a large number of photoreceptor cells from microscopy images of whole retinas. We use the data and the model to study what exactly it means for a cell to have a binary or graded cell fate and how these cell fates can be distinguished from each other. Our study shows how tens-of-thousands of individual photoreceptor cells can be patterned across a complex tissue by a regulatory network, creating a different outcome depending upon the received inputs.

## Introduction

How the numerous neuronal subtypes of the vertebrate nervous system are patterned is an ongoing puzzle in developmental neurobiology. Are neuronal subtypes distinct states generated by binary gene expression decisions? Or are they highly complex with ranges of graded gene expression? The answers likely lie somewhere in between, with some genes expressed in a simple switch-like fashion and other genes expressed across a range of levels to define cell fate. A challenge is to understand how cells interpret regulatory inputs to generate complex patterns of binary and graded cell fates across tissues. Here, we address this question in the context of cone photoreceptor patterning in the mouse retina.

Photoreceptors detect and translate light information into electrical signals, triggering the neuronal network yielding visual perception. There are two main classes of image-forming photoreceptors: rods and cones. Rods are mainly used in night vision, while cones are used in daytime and color vision. In most mammals, cones express S-opsin, which is sensitive to blue or UV-light, and M-opsin, which is sensitive to green light (Calderone and Jacobs, 1995; Wang et al., 2011b).

The common laboratory mouse, *Mus musculus*, displays complex patterning of cone opsin expression across its retina, providing an excellent system to study binary and graded features of cell fate specification. The dorsal third of the retina is mostly comprised of cones that express M-opsin, and a minority that exclusively express S-opsin. In the central region, most cones co-express S- and M-opsin, with small subsets that express only S- or only M-opsin. The majority of the ventral region contains cones that co-express S- and M-opsin, with significantly higher levels of S-opsin compared to M-opsin (Applebury et al., 2000; Baden et al., 2013; Calderone and Jacobs, 1995; Haverkamp et al., 2005; Rohlich et al., 1994; Szel et al., 1994). Here, we expand upon these pioneering studies to examine cone patterning along the complete dorsal to ventral (D-V) axis of the mouse retina and quantitatively model how regulatory inputs influence cone cell patterning.

Cone subtype fate is not only characterized by opsin expression, but also connectivity. Two cone subtypes have been defined primarily on connectivity to downstream bipolar neurons. 3-5% of cones are “genuine” S cones that express S-opsin only and connect to blue-cone bipolars. The remaining cones express S-opsin only, M-opsin, or both S-opsin and M-opsin and do not connect to blue-cone bipolars (Haverkamp et al., 2005). The regulatory relationship between connectivity and opsin expression during cone subtype specification has not been established. In this work, we focus on the binary and graded nature of opsin expression, specifically examining this aspect of cone subtype fate.

Cone opsin expression is regulated by thyroid hormone (TH) signaling. TH and the nuclear thyroid hormone receptor Thrβ2 are important for activating M-opsin and repressing S-opsin expression (Roberts et al., 2006). TH exists in two main forms: T4, the circulating form, and T3, the form that binds with high affinity to nuclear receptors and acts locally to control gene expression (Samuels et al., 1974; Schroeder et al., 2014). T3 levels are highest in the dorsal part of the mouse retina and decrease ventrally (Roberts et al., 2006). Deiodinase 2 (Dio2), an enzyme that converts T4 to T3, is expressed at high levels in the dorsal region of the mouse retina, and is thought to maintain the gradient of T3 in the adult retina (Bedolla and Torre, 2011; Corbo et al., 2007). T3 is sufficient to induce M-opsin expression and repress S-opsin expression (Roberts et al., 2006).

Thrβ2, a receptor for TH, is expressed in all cones of the retina (Roberts et al., 2005; Sjoberg et al., 1992). Thrβ2 acts as a transcriptional repressor in the absence of T3 binding, and as a transcriptional activator when T3 is bound (Bernal, 2005). Thrβ2 activity is required for expression of M-opsin and repression of S-opsin (Applebury et al., 2007; Eldred et al., 2018; Ng et al., 2001; Pessoa et al., 2008; Roberts et al., 2006; Suzuki et al., 2013). Additionally, RXRγ, a hetero-binding partner of Thrβ2, is necessary for repressing S-opsin in dorsal cones (Roberts et al., 2005). The transcription factors Vax2 and Coup-TFII, which regulate and respond to retinoic acid levels, have also been implicated in photoreceptor patterning (Alfano et al., 2011; Satoh et al., 2009). For this study, we focus on modeling the contributions of TH and Thrβ2 to cell fate outcomes.

We desired to quantitatively model cone fate specification in the mouse retina. Our current theoretical understanding of cell fate determination within a tissue describes individual cell types as distinct valleys on an “epigenetic landscape” (Furusawa and Kaneko, 2012; Micheelsen et al., 2010; Wang et al., 2011a; Zhang and Wolynes, 2014; Zhou et al., 2012). Cells make fate decisions by transitioning to one of these “attractor” states on the landscape (Olsson et al., 2016). Differences in gene expression between the states give rise to phenotypic differences between cell types. However, clustering based on single-cell transcriptomics data alone misses subpopulations unless hidden variables are accounted for (Buettner et al., 2015; Setty et al., 2019).

Recently, computational work has also focused on developing mechanistic models of cell-fate decisions (Olariu and Peterson, 2019; Rothenberg, 2019; Teles et al., 2013), especially the formation of patterns in time and space (Formosa-Jordan, 2018; Liang et al., 2015). Multiscale approaches that combine stochastic and deterministic models of tissues at the scale of individual cells have shown promise in helping to elucidate the details of tissue patterning (Coulier and Hellander, 2018; Engblom, 2018; Engblom et al., 2018; Folguera-Blasco et al., 2019; Johnston et al., 2011). The zebrafish retina has been studied to model cell fate decision making based on anticlustering mechanisms that give rise to a lattice structure of differentiated cell types (Cameron and Carney, 2004; Ogawa et al., 2017; Tyler et al., 2005). The near perfect grid of cone subtypes in the zebrafish eye is in stark contrast with the variable patterning of cone subtypes in the D-V axis of the mouse retina (Baden et al., 2013; Haverkamp et al., 2005; Viets et al., 2016).

Here, we present a multiscale model describing the emergence of the complex arrangement of cone cells found in the mouse retina using both stochastic and deterministic methods. We collect data for the model from analysis of immunofluorescence images of retina tissues to identify and map individual cones along the entire D-V axis of the mouse retina. Based on opsin expression in the individual cells, we find that cones can be classified into two main subtypes: S-only cones and co-expression competent (CEC) cones. The S-only cones express S-opsin only, whereas the CEC cones express M- and/or S-opsins in opposing dorsal-ventral gradients, with higher levels of M-opsin in cones in the dorsal retina and higher levels of S-opsin in cones in the ventral retina. We then use the data to parameterize a mathematical model of a two-step cone patterning process. Step one is a binary choice between S-only fate and CEC fate. If CEC fate is selected, a second mechanism regulates S- and M-opsin expression in a reciprocal, graded manner, along the dorsal-ventral axis. Our quantitative modeling shows that the expression of S- and M-opsins in CEC cells are differentially activated based on dorsal-ventral patterning inputs from T3. Our model closely recapitulates cone patterning observed in the mouse retina and provides insights into how spatial patterning inputs regulate binary and graded features of cell fate in parallel.

## Results

### Characterization of cone subtype patterning in the mouse retina

To globally characterize patterning of opsin expression in the mouse retina, we first examined the relative intensity of S- and M-opsin expression in the D-V and temporal-nasal (T-N) axes at low resolution for whole-mounted retinas. We immunostained and imaged S- and M-opsin proteins at 100X magnification (see Materials and Methods). Following image acquisition, we manually rotated each image so that the D-V axis was aligned vertically (**Fig. 1A, E, I)**. At this resolution, individual cells cannot be identified, so we instead subdivided each image using a 25 pixel × 25 pixel grid, which is an area containing approximately one to two cells. Within each bin of the grid, we counted the number of pixels that had significant S-opsin signal alone, M-opsin signal alone, or both M- and S-opsin signals. We then normalized each bin by the total number of pixels with expression in that bin. This calculation gave us the relative density of each photoreceptor type by location in the retina (**Fig. 1B, F, J)**.

**Figure 1.**
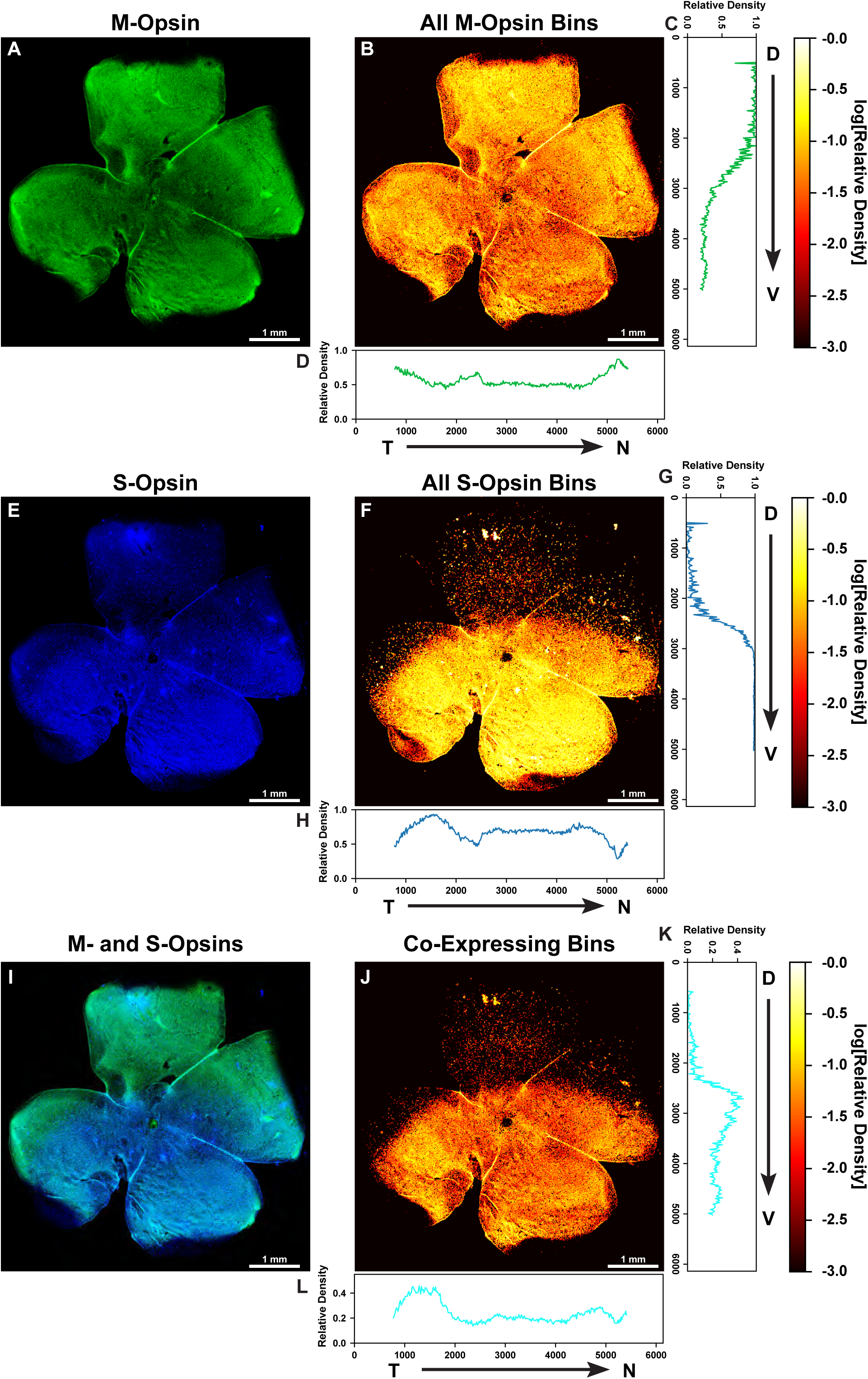
Analysis of opsin expression intensity across the mouse retina. **A, E, I)** Whole mounted C57BL/6 mouse retina stained for M-opsin (green) and S-opsin (blue). **B, F, J)** Heatmap displaying the log relative density of pixels that have opsin signal identified in a 25 mm**^2^** region. **A)** M-opsin signal. **B)** Heatmap of total M-opsin density bins. **C)** Graph of the relative density of pixels that are expressing M-opsin summed horizontally (D – V). **D)** Graph of the relative density of pixels that are expressing M-opsin summed vertically (T – N). **E)** S-opsin signal. **F)** Heatmap of total S-opsin density bins. **G)** Graph of the relative density of pixels that are expressing S-opsin summed horizontally (D – V). **H)** Graph of the relative density of pixels that are expressing S-opsin summed vertically (T – N). **I)** M-opsin and S-opsin (co-expression) signal. **J)** Heatmap of co-expressing opsin density bins. **K)** Graph of the relative density of pixels that are co-expressing S- and M-opsin summed horizontally (D – V). **L)** Graph of the relative density of pixels that are co-expressing S- and M-opsin summed vertically (T – N). T = Temporal, N = Nasal, D = Dorsal, V = Ventral.

Next, we quantified global differences in patterning in the D-V and temporal-to-nasal (T-N) dimensions. We averaged the binned density values to obtain the relative density as a function of either D-V (**Fig. 1C, G, K**) or T-N position (**Fig. 1D, H, L**). We observed distinct transitions in both S- and M-opsin expression along the D-V axis. High levels of M-opsin in the dorsal region exhibit a gradual transition to low levels in the ventral region (**Fig. 1C**). In contrast, S-opsin shows a rapid transition from zero to high expression in the D-V axis (**Fig. 1G**). As these opsins display an inverse yet non-complementary relationship, co-expression was most prominent in the middle third of the retina where these two transitions overlap (**Fig. 1K**). We observed minimal variation in S- and M-opsin signal in the T-N axis (**Fig. 1D, H, L**). We imaged and analyzed six wild-type retinas at this resolution and saw a similar pattern in each (Table S1). Together, we observed differential graded patterning for S- and M-opsin expression along the D-V axis (**Fig 1C, G, K**).

### Analysis at single cell resolution reveals two distinct cone subtype populations

To further investigate photoreceptor patterning along the D-V axis, we next analyzed cone subtype specification at the single-cell level. For the same six retinas, we imaged S- and M-opsin expression at 200X magnification in a strip measuring approximately 600 μm × 6,000 μm aligned vertically along the D-V axis (**Fig. 2A**). Previous studies analyzed ∼500 μm in the dorsal-ventral axis centered on the transition region (Haverkamp et al., 2005), whereas our approach enabled evaluation of the entire ∼6,000 μm length of the retina.

**Figure 2.**
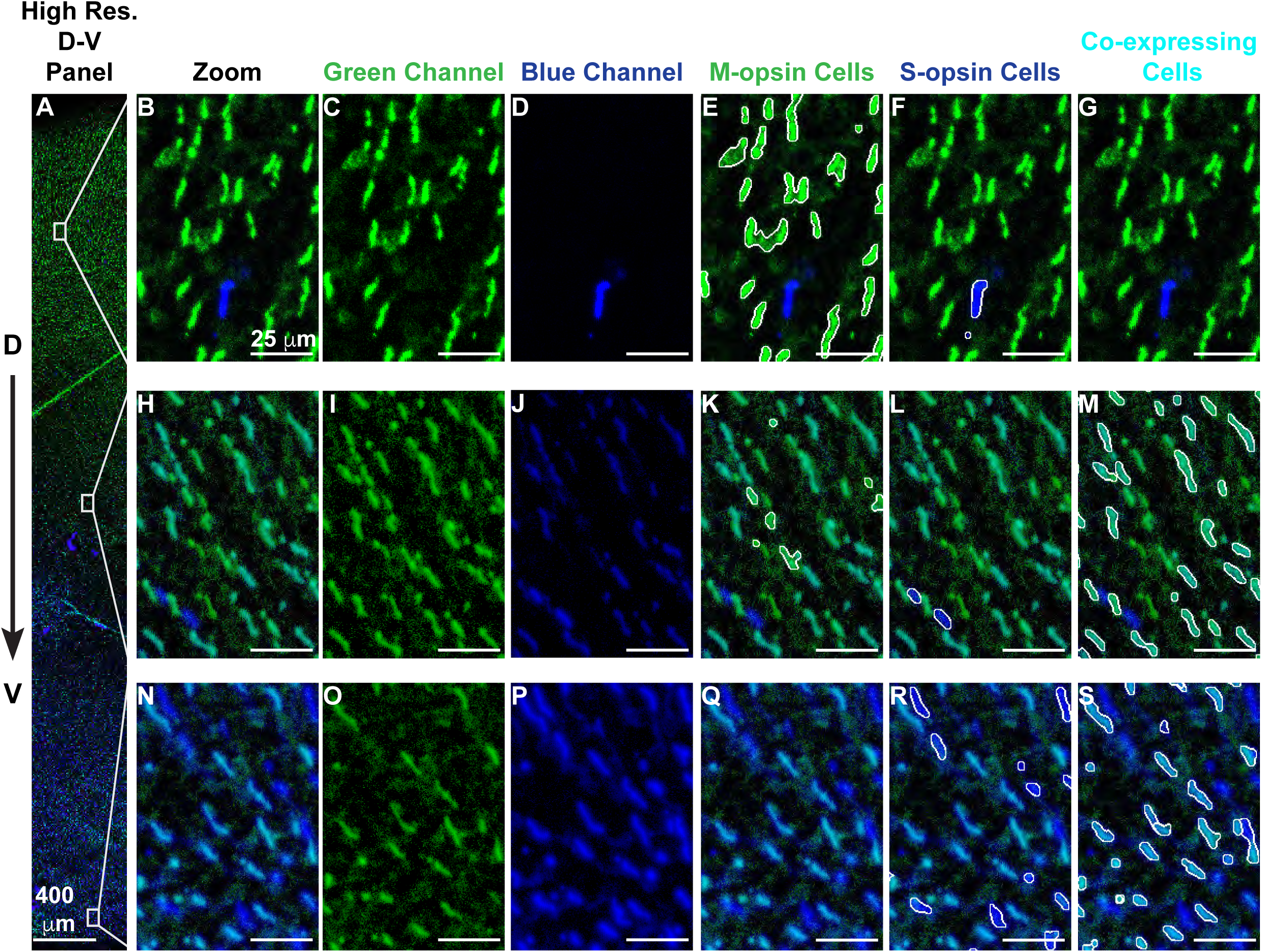
Identification of cone subtypes. **A-S)** Retina stained with antibodies against M-opsin (green) and S-opsin (blue) **A)** High-resolution region spanning the dorsal to ventral retina. **B-G)** A region of the dorsal retina. **F-M)** A region of the central retina. **N-S)** A region of the ventral retina. **B, H, N)** Blue and green channels. **C, I, O)** Green channel only. **D, J, P)** Blue channel only. **E, K, Q)** White outline indicates identified M-opsin expressing cells. **F, L, R)** White outline indicates identified S-opsin expressing cells. **G, M, S)** White outline indicates identified co-expressing cells.

At 200x magnification, we were able to distinguish and identify individual cells. We developed an analysis pipeline to identify the position, size, and boundaries of each cell, a process known as segmentation (see **SI Methods, 1.1.1**). Overall, we identified ∼250,000 total cells across six retinas. Using these cell boundaries, we calculated the expression intensity of M- and S-opsin for each cell. We classified cones into groups expressing M-opsin only (**Fig. 2E, K, Q**), S-opsin only (**Fig. 2F, L, R**), and S- and M-opsin co-expression (**Fig. 2G, M, S**). The pipeline did not identify distinct morphological or size differences among the cones (data not shown). We found that the pipeline’s accuracy and false positive rate were comparable to hand-scored retinas (see **SI Methods 1.1.2**).

After obtaining the cell boundaries, we quantified the density of cone subtypes based on opsin expression relative to D-V position. Consistent with our low-resolution analysis, we observed a gradual decrease in the abundance of cones expressing M-opsin in the dorsal to ventral direction (**Fig. 3A**), contrasted by a sharp increase in S-opsin expressing cones (**Fig. 3B**). We fit these curves to Hill functions to quantify the steepness of the transition (**Fig. S1**). The transition in the S-opsin expressing cells is extremely sharp with an average Hill coefficient of ∼30 while the M-opsin transition is much more gradual with a coefficient of ∼2-3.

**Figure 3.**
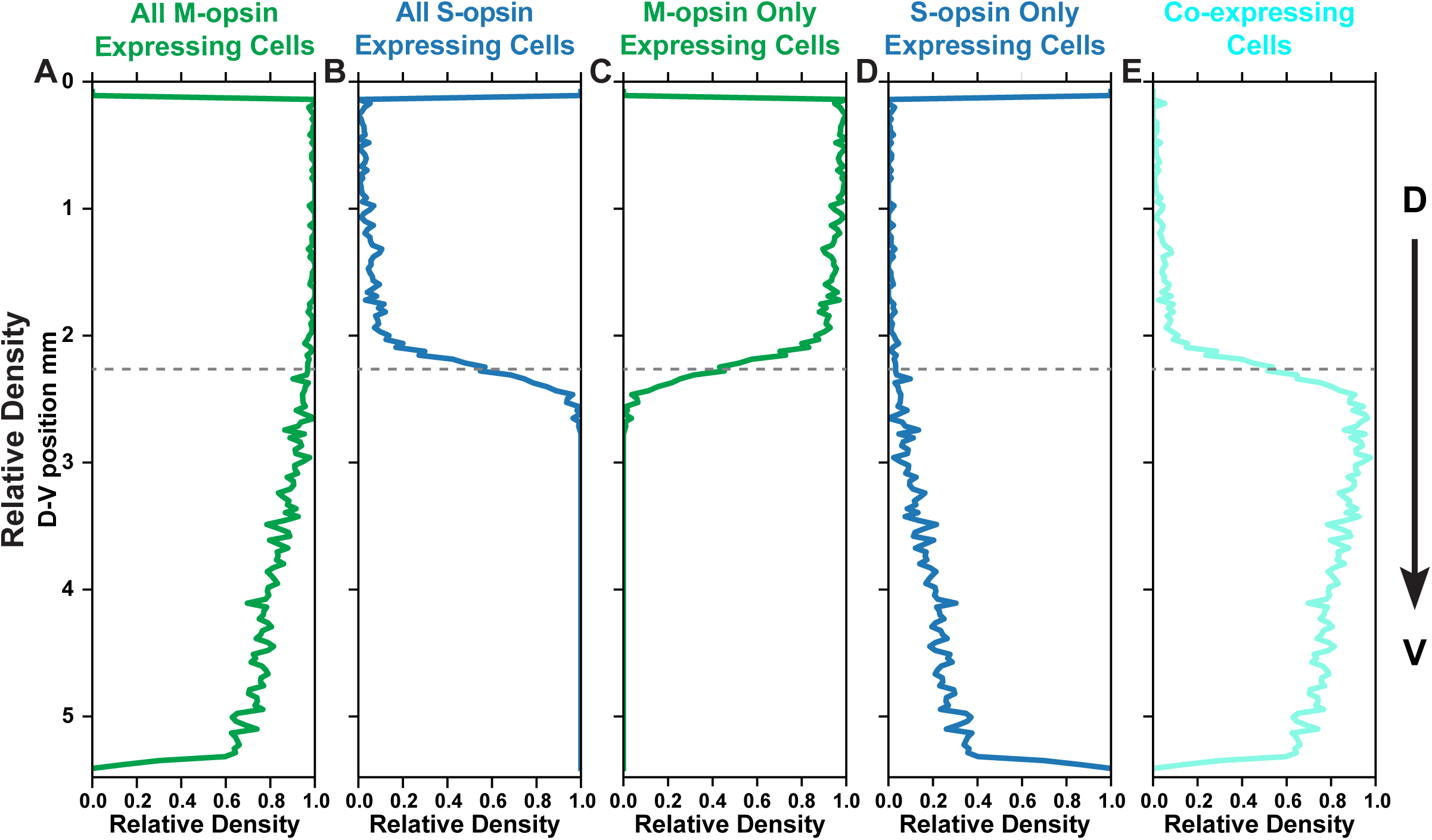
Spatial distribution of M- and S-opsins in cone cells. Relative density of a cone population summed horizontally across the image and displayed in the dorsal to ventral position. Dotted line represents midpoint of transition zone. **A)** All M-opsin expressing cells. **B)** All S-opsin expressing cells. **C)** M-opsin only expressing cells. **D)** S-opsin only expressing cells. **E)** Co-expressing cells.

To compare the transition region between retinas, we established a reference point to align the images. Since the S-opsin transition is sharp and an external reference is absent, we used the midpoint of the S-opsin transition from the fit as the reference point. We aligned all of the retinas and overlaid the transition fits (**Fig. S2**). The relative position of the S-opsin and M-opsin transitions are consistent from retina to retina, suggesting that the transitions in S-opsin and M-opsin expression are driven by a common effector. M-opsin only expressing cones decline at the transition point (**Fig. 3C**), coincident with the dramatic increase in S-opsin expression (**Fig. 3B**). At this transition point, cones begin to express both S- and M-opsins (**Fig. 3E**) and the cone populations are very diverse, comprised of those expressing M-only, S-only, and varying levels of both S- and M-opsins (**Fig. 2H-M**). The fraction of S-only cells gradually increases from ∼1% of cones in the dorsal region to ∼20-30% in the ventral region (**Fig. 3D, S3**). These analyses show the differential, inverse responses of S- and M-opsin expression to D-V patterning inputs on the individual cell level.

In a previous study, Haverkamp et al. measured differential opsin expression of cone cells in a window of ∼500 μm near the transition point (Haverkamp et al., 2005). In agreement with our data, they observed that ∼8-20% of cones expressed only S-opsin. Our results show that this measurement was part of a broader binary decision trend extending much further along the D-V axis in both directions. They also discovered that within this population, in the ventral region where S-only cones are more abundant, about 5% of S-opsin only cones contact S-cone bipolar cells and they classified these as genuine S-cones. These genuine S-cones are evenly distributed across the retina (Haverkamp et al., 2005).

To see if we could distinguish these two classes of cone subtypes in our data, we calculated the joint probability distribution for S- and M-opsin intensity in individual cells for each retina (**Fig. 4A, S4**). When considering all cones in the retina, there appear to be three clusters of cell-types corresponding to the three classifications that we defined earlier: S-only, M-only, and co-expressing. However, when the data are analyzed by D-V position (**Fig. 4B-F, S5**), a different pattern emerges. First, we see an S-only cluster than has high and consistent expression of S-opsin while increasing in abundance along the D-V axis (**Fig. 4**). Interestingly, the other two clusters resolve into a single cluster that changes position in a continuous way, gradually moving from low S-opsin expression and high M-opsin expression in the dorsal region, to moderate S-opsin expression and low M-opsin expression in the ventral region (**Fig. 4**).

**Figure 4.**
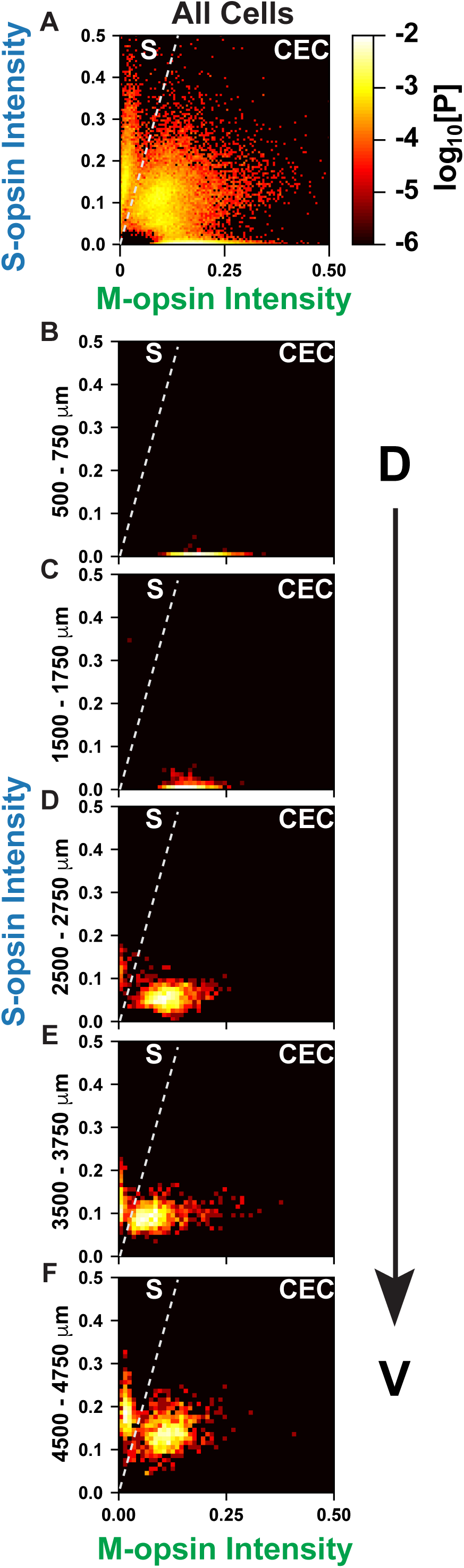
S- and M-opsin intensities in cones. Cones are ranked according to the intensity of S- and M-opsin expression levels. Intensity values are represented in arbitrary units. Each point is colored according to the log10[Probability] of expression levels. A line is drawn on the graph to show the separation between the two discrete populations of S-opsin only and CEC cone populations. **A)** All cones in the regions imaged. **B)** Cones in the dorsal 500-750 mm. **C)** Cones in the dorsal 1500-1750 mm. **D)** Cones in the central 2500-2750 mm. **E)** Cones in the ventral 3500-3750 mm. **F)** Cones in the ventral 4500-4750 mm.

Thus, these data suggest that the mouse retina contains two main subtypes of cones: 1. S-only cones that have high S-opsin expression independent of D-V position and 2. co-expression competent (CEC) cones that express S- and/or M-opsins dependent upon D-V position. In the ventral region there is a mixture of S-only cones and CEC cones that express M-opsin at a very low level. It is difficult to distinguish these two classes using only S-opsin expression, but as can be seen in **Fig 4**., the two populations are well separated when comparing both M- and S-opsin intensities. The S-only cones identified with our approach may, therefore, contain a subclass corresponding to the genuine S-cones identified by Haverkamp et al., but as we could not distinguish their connectivity to bipolar cells, we are only able to describe the populations of S-opsin only expressing cones.

### Expression levels of S- and M-opsin in cone cell subtypes

Having classified the major subtypes of cones and related their positions and opsin expression states, we next evaluated the D-V dependence of the opsin expression intensity in individual cones. We quantified opsin expression for all M-opsin expressing cones (**Fig. 5A, F**), all S-opsin expressing cones (**Fig. 5B, G**), M- and S-opsin CEC cones (**Fig. 5C, H**), M-opsin only CEC cones (**Fig. 5D, I**), and S-opsin only cones (**Fig. 5E, J**) relative to their D-V positions within the retina.

**Figure 5.**
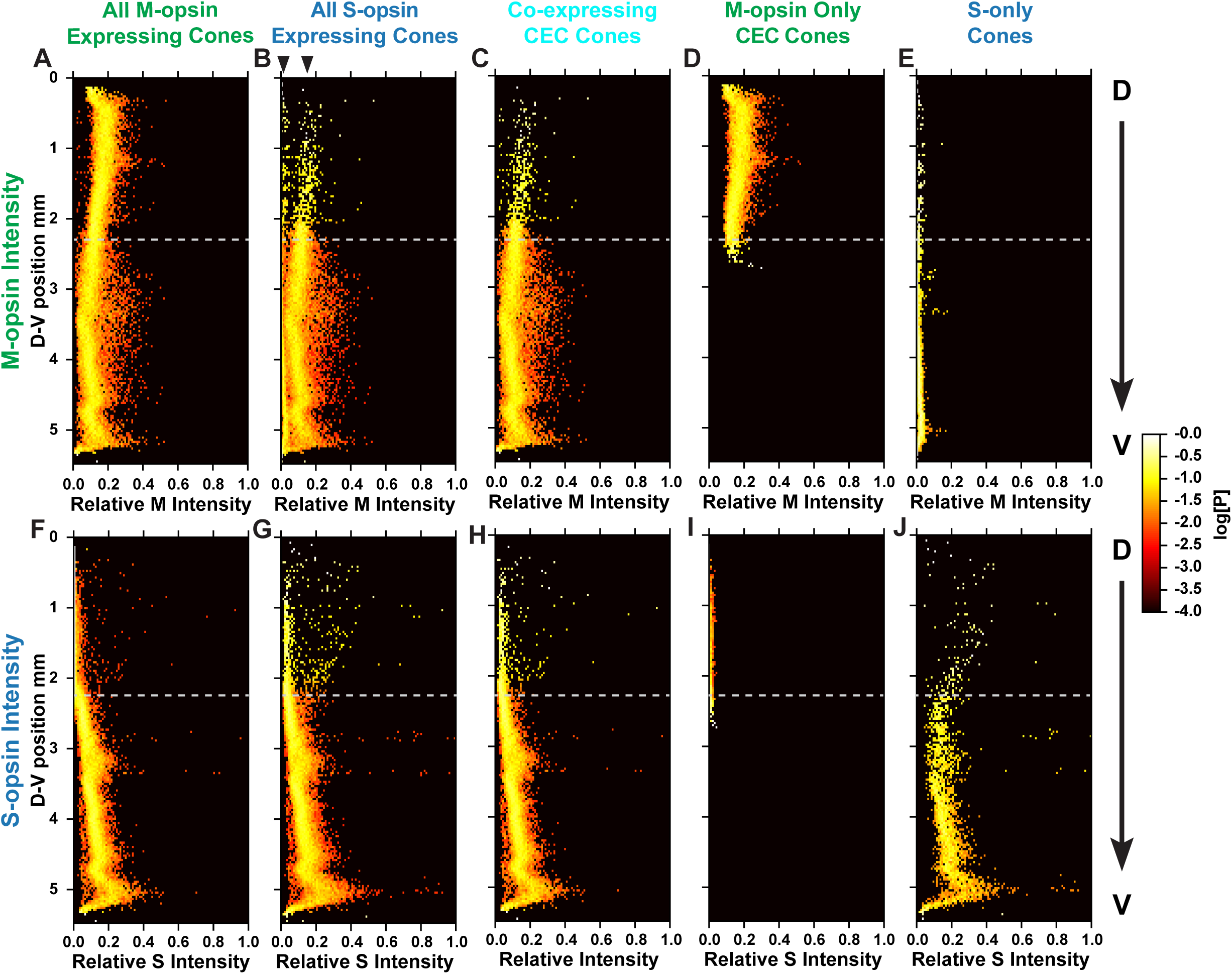
Intensity of M- and S-opsins in cones. Relative intensity of M- or S-opsin in a cone population (X-axis) is displayed as a function of dorsal to ventral position. Each point is colored according to the log10[Probability] of expression levels. **A-E)** Relative intensity of M-opsin expression **F-J)** Relative intensity of S-opsin expression **A, F)** All M-opsin expressing cells. **B, G)** All S-opsin expressing cells. For **B)**, arrow heads mark two distinct groups of cells in the dorsal region. **C, H)** CEC cones co-expressing both S- and M-opsins. **D, I)** M-opsin only expressing CEC cones. **E, J)** S-opsin only expressing CEC cones.

In CEC cones, M-opsin expression levels decrease in the D-V axis, with the midpoint of expression level located at the transition point (**Fig. 5A, C, D, S6**). In contrast, S-opsin expression in CEC cells is very low in the dorsal region and increases linearly in the D-V axis starting at the transition point (**Fig. 5F, H, I, S6**). The slope of increase for S-opsin is steeper than for the M-opsin decrease (**Fig. S7**).

Compared to CEC cones, S-only cones have an overall higher expression level of S-opsin, particularly in the dorsal region (**Fig. 5J** compared to **G** and **H**). M-opsin expression in S-only cones is significantly lower than the lowest M-opsin expression seen in CEC cones (**Fig. 5E**). In **Fig. 5B**, this difference can be seen as two distinct lines of density (**Fig. 5B**, arrow heads). Together, these analyses defined the expression of S- and M-opsin in the two cone populations in relation to their D-V positions in the retina.

### Modeling cone subtype fate decisions

To interrogate how regulatory inputs could produce the complex pattern of binary and graded cell fates in the mouse retina, we developed a multiscale model describing the probability distributions of the cone subtype decisions (i.e. binary choice) and S- and M-opsin expression levels (i.e. graded) as functions of position along the D-V axis (**Fig. 6, SI Methods 1.2**). We modeled a 5 mm × 1 mm × 5 µm section of the retina with the long dimension aligned with the D-V axis.

**Figure 6.**
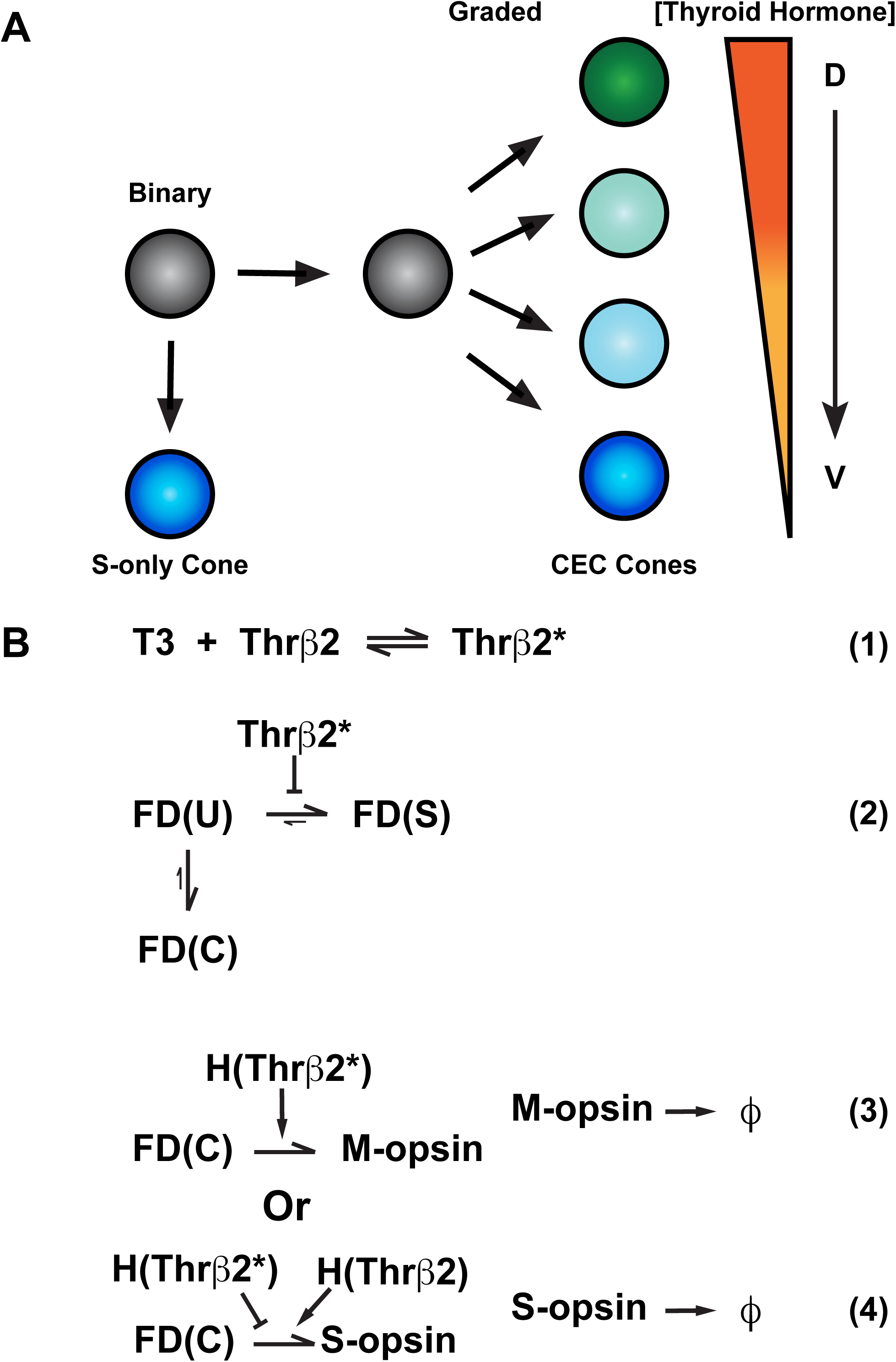
Model for cone cell fate specification. **A)** A naïve cell (grey) makes a binary decision between S-opsin only (blue) or co-expressing competent (CEC) cone fate (green, cyan or blue). The CEC cone expresses graded levels of M- and S-opsin dependendent on the D-V concentration of thyroid hormone. **B1-4)** T3 (Thyroid hormone), Thrβ2* (active Thrβ2 binding T3), FD (fate determinate function), U (undifferentiated cell), S (S-only cone), C (Co-expressing cone), H (Hill function), ϕ (degradation constant of opsin proteins). **B1)** Binding of T3 to Thrβ2 activates Thrβ2 (Thrβ2*) **B2)** Thrβ2 controls the binary decision between S-opsin only/FD(S) or CEC/FD(C) cone fate **B3)** Thrβ2* promotes M-opsin expression **B4)** Thrβ2* inhibits S-opsin expression, whereas inactive Thrβ2 promotes S-opsin expression

TH signaling activates M-opsin expression and represses S-opsin expression (Ng et al., 2001; Roberts et al., 2006). T3 is a critical regulator of cone subtype fate in the human retina (Eldred et al., 2018), and scRNA-seq data suggest that Thrβ2 is expressed in all mouse cones (Clark et al., 2019). Though other diffusible factors and transcription factors play roles (Alfano et al., 2011; Roberts et al., 2005; Satoh et al., 2009), TH signaling is the main and best-understood determinant of cone subtype fate. Thus, we built a simplified model of cone subtype specification based on the dorsal-ventral regulation of cone fates by the gradient of T3.

Within the modeled volume, T3 molecules diffuse according to the deterministic diffusion equation with constant concentration boundaries, establishing a D-V gradient (**Eq. S1, SI Methods**). Also, within the volume, we stochastically modeled ∼23,000 individual cones spaced on a hexagonal grid (**Fig. 7**). These cells stochastically exchange T3 molecules with the surrounding deterministic microenvironment. Within each cone, T3 can bind to and activate Thrβ2 (Thrβ2*), controlling both fate specification and opsin expression (**Fig. 6B1-4**).

**Figure 7.**
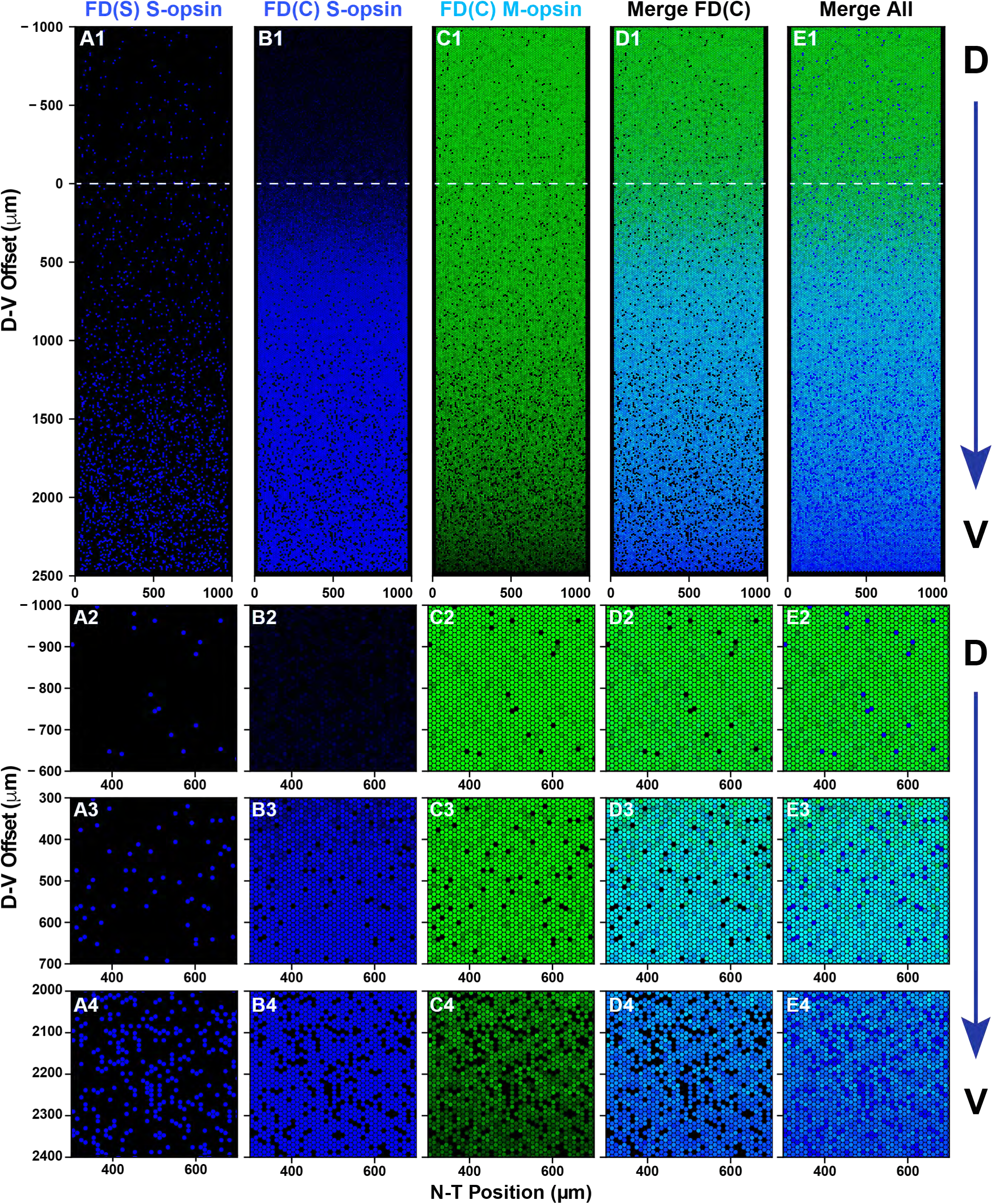
Simulated cone mosaic produced by the quantitative model. Simulated cone photoreceptor mosaic generated by the quantitative model displaying expression of S-opsin (blue), and M-opsin (green). A dorsal to ventral region is shown. **A1, B1, C1, D1)** Complete simulated D-V strip. **A2, B2, C2, D2)** Zoom in the dorsal region. **A3, B3, C3, D3)** Zoom in the central region. **A3, B3, C3, D3)** Zoom in the ventral region. **A1-4)** S-opsin only cones. **B1-4)** S-opsin expression in CEC cones. **C1-4)** M-opsin expression in CEC cones. **D1-4)** S- and M-opsin expression in CEC cones **E1-4)** All cones including S-opsin only and CEC cones.

For the binary fate decision, we defined a fate determinant function, FD(X). Photoreceptors start in an undifferentiated fate, FD(U), and stochastically progress to either the FD(S) (S-only) fate or FD(C) (CEC) fate (**Fig. 6B2**). Selection of the FD(S) fate is negatively influenced by Thrβ2* (**Fig. 6B2**). Once cells enter the FD(S) fate, S-opsin is constitutively expressed at a high level regardless of D-V position (**Fig. 6B2**). In FD(C) cones, M-opsin expression is induced by Thrβ2* (**Fig. 6B3**). Conversely, S-opsin in FD(C) cells is negatively regulated by Thrβ2* and positively regulated by inactive Thrβ2 receptors (**Fig. 6B4**). Full details of the model are given in the SI Methods 1.2, along with parameterization details.

### Model-based simulations recapitulate experimental cone patterning

To compare the output of our probabilistic model to experimental data sets, we ran a set of 100 individual simulations and calculated the probability distributions of various observables. **Fig. 7** shows the output of one simulation. Moving from dorsal to ventral, the model reproduces the gradual increase in the fraction of S-only cones, FD(S) (**Fig. 7A1-4**), as well as the sharp transition in CEC cones, FD(C), expressing S-opsin at the transition zone (**Fig. 7B1-4**). Similarly, we observed the gradual decrease in the fraction of CEC cones expressing M-opsin (**Fig. 7C1-4**). In the overlapping region, there are a significant number of cones that co-express both S- and M-opsins (**Fig. 7D1-4**). In the dorsal region, a small number of S-only cones that highly express S-opsin are readily apparent (**Fig. 7A1-2, E1-2**).

To characterize how well our model recapitulated the observed experimental cell distributions, we calculated the mean density of cells of various phenotypes as a function of D-V position (**Fig. S8)**. These average density profiles compare well to the experimental density profiles shown in **Fig. 3**. Together, cone fate patterning and expression levels are highly similar in our model and the imaged retinas: S-only cells (**Fig. 7A2-4** compared to **Fig. 2F, L, R**), S-opsin expression in CEC cones (**Fig. 7B2-4** compared to **Fig. 2E, G, K, M, Q, S**), and M-opsin expression in CEC cones (**Fig. 7C2-4** compared to **Fig. 2E, G, K, M, Q, S**).

We parameterized our model using the mean of all the retinas sampled, which exhibited retina-to-retina variability (**Fig. S1, S3, S7**). Therefore, it is not expected that our model will exactly recapitulate the patterning of any individual retina.

We next calculated the probability distributions of S- and M-opsin expression along the D-V axis for our simulation data (**Fig. S9**). The mean intensity of M-opsin in CEC cones gradually decreases as D-V position increases. The S-opsin distribution shows high expression in the ventral-most region, but has two separate populations in the dorsal region: the highly expressing S-only cells and the lowly expressing CEC cells. The CEC cones converge to zero S-opsin expression in the dorsal region while the S-only cones maintain high expression as they decrease in abundance. Because our simulated distributions are constructed from 100 independent simulations, the probability density of the S-only cones is much smoother than in the experimental data (compare **Fig. S6 and S9**). The simulated expression features are in agreement with the experimental expression profile (**Fig. 5, S6**).

We next related the joint probability distributions for the experimental (**Fig. 8A-E**) and simulated (**Fig. 8F-J**) data along the D-V axis. The simulated and experimental data show two distinct populations: 1) S-only cones with high S-opsin expression and no M-opsin expression whose expression levels are independent of D-V position, and 2) CEC cones that gradually change from high M-opsin and low S-opsin expression to moderate M-opsin and high S-opsin expression along the D-V axis. **Fig. S10** shows the joint probability distribution between S- and M-opsin expression within 250 µm D-V bins for all 100 simulations. In the high resolution simulated data, it is evident that the position of the CEC cell cluster gradually changes with D-V position. Our model closely simulates the experimental data, and supports the hypothesis that cells respond differentially to the same morphogen gradient, producing both binary and graded cell fates.

**Figure 8.**
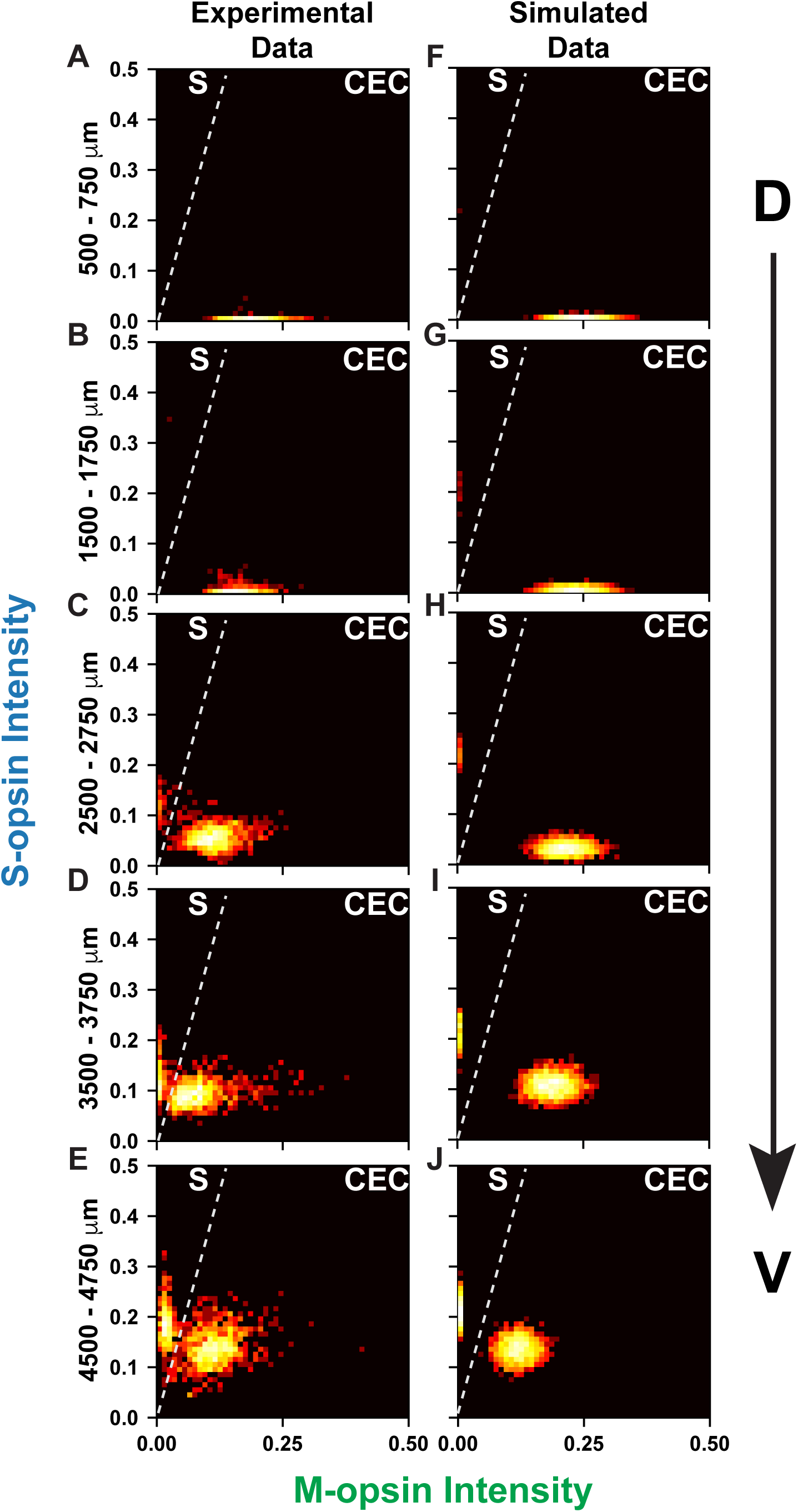
D-V cone pattering in simulated and experimental data. Cones are ranked according to the intensity of S- and M-opsin expression levels. Intensity values are represented in arbitrary units. Each point is colored according to the log10[Probability] of expression levels. **A-E)** Experimental data, as seen in Fig. 4B-F. **F-J)** Simulated data. **A, F)** Cones in the dorsal 500-750 mm. **B, G)** Cones in the dorsal 1500-1750 mm. **C, H)** Cones in the central 2500-2750 mm. **D, I)** Cones in the ventral 3500-3750 mm. **E, J)** Cones in the ventral 4500-4750 mm.

### Correlation between S-opsin and CEC fate decisions

Our quantitative simulations give us the capacity to test various hypotheses about retinal patterning. We wanted to know whether the gradual decrease in the CEC cone population and the sharp increase in S-opsin expression in these CEC cells were driven by a shared upstream signaling input. If these two processes respond to the same upstream input, we would expect that they should be coupled and be linked by D-V position. If, however, they do not respond to the same input we would expect that their transitions should be independent. In our model, they are coupled through the T3 gradient and we wanted to test if the experimental retinas displayed more or less variability than expected. Because stochasticity in both the experimental data and our model introduces retina-to-retina variability, we checked for overlap of the corresponding probability distributions.

We calculated the probability of having a given CEC population fraction at the S-opsin transition point from both our simulations and experimental retinas (**Fig. 9A**). We also calculated the rate at which the CEC population decreases at the transition point (**Fig. 9B**). As can be seen from both plots, the two probability distributions are in good agreement. In particular, the widths of the experimental probability distributions are similar to the widths from the stochastic simulations. With the small number of experimental data points, we do not assign a level of statistical significance to the overlaps, but they provide qualitative evidence that the two transitions are in fact coupled through a shared upstream input.

**Figure 9.**
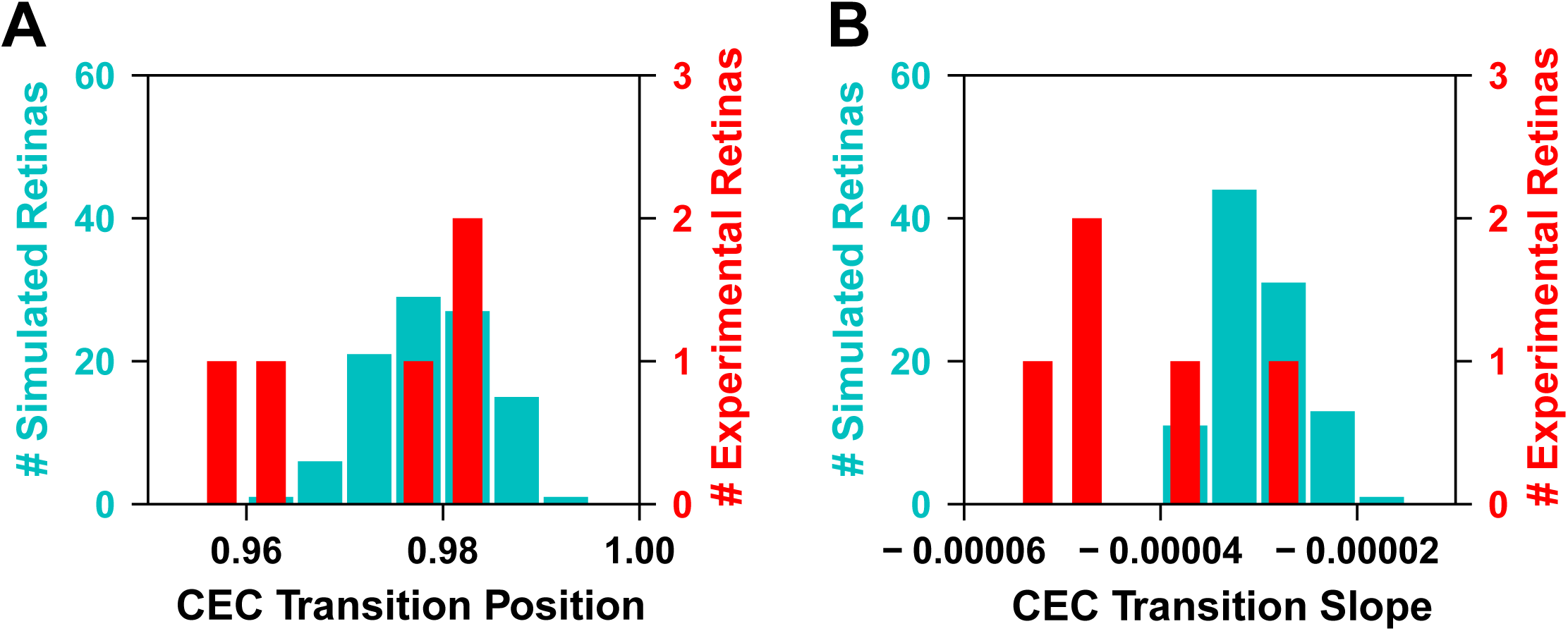
Correlation between CEC fate and S-opsin transitions. **A)** The fraction of CEC cells at the point where the S-opsin transition is at its midpoint. Data are shown for both experimental (red) and modeled (cyan) retinas. **B)** The slope of the CEC transition at the S-opsin midpoint, for both experimental (red) and modeled (cyan) retinas. Note: the distributions of only 5 of the 6 retinas are included here, as one of the images had major disruptions at the transition zone due to disecting and mounting.

### Comparison of Thrβ2 mutant retinas to wild-type retinas

Finally, to elucidate the effect of the T3 gradient on cone cell patterning in the mouse retina, we dissected and imaged Thrβ2 knockout mutant retinas (ΔThrβ2). In the absence of functional Thrβ2, no M-opsin is expressed (Applebury et al., 2000; Ng et al., 2001; Roberts et al., 2006). Consistent with previous work, we observed no M-opsin expression in these mutant retinas. Fluorescence from anti-M-opsin antibodies was nonspecific and stained cell and background with equal intensity (**Fig. S11**).

The expression of S-opsin in cones was also markedly different between ΔThrβ2 and WT retinas. Both the density of S-opsin-expressing cones and the expression distribution are flat with respect to the D-V axis (**Fig. 10B, S12**). Also, the relative intensity of the S-opsin signal across the retina was much lower in ΔThrβ2 retinas than the maximum value seen in WT retinas (e.g., from cones in the ventral region). ΔThrβ2 retinas and wildtype retinas were taken at the same time and stained with the same batch of antibody, then imaged with the same laser intensity for comparison of opsin levels. We found that the relative intensity of opsin staining in cones was most similar to the middle region of WT retinas. This effect is consistent with our model in which S-opsin expression is controlled through a combination of negative regulation by active Thrβ2* and positive regulation by inactive Thrβ2 (**Fig. 6B4**).

**Figure 10.**
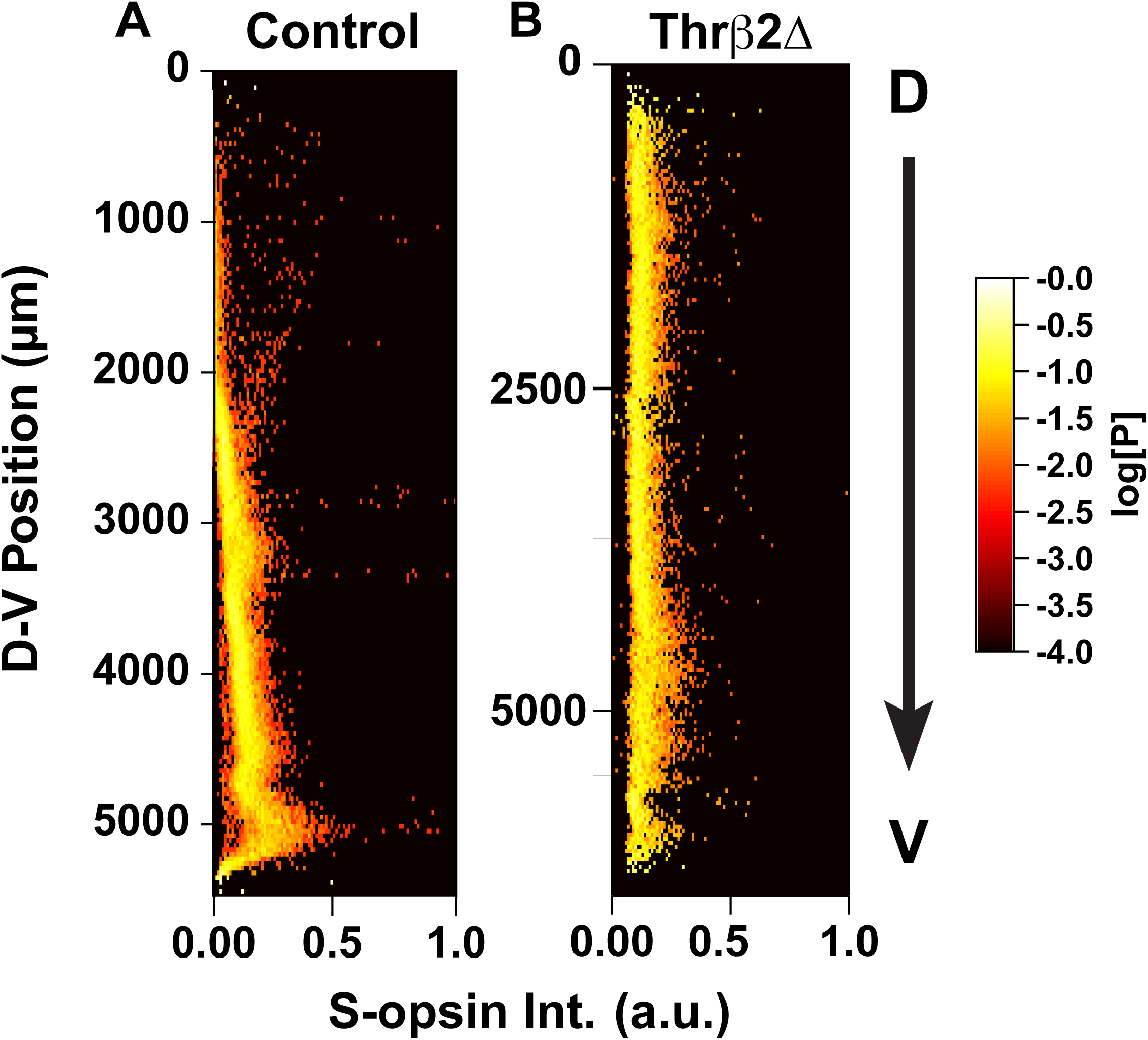
ThrB2Δ mouse Intensity Plots. Relative intensity of S-opsin cone cells (X-axis) displayed as a function of dorsal to ventral position. Each point is colored according to the log10[Probability] of expression levels. **A)** Control Retina, as seen in Fig. 5F. **B)** ThrB2Δ retina.

## Discussion

In these studies, we described the distribution of cone photoreceptors in the mouse retina and developed a quantitative model for the specification of binary and graded cell fates in response to D-V regulatory inputs. By using high-resolution microscopy combined with automated image analysis, we expanded on previous studies and mapped the cell fate decisions of cone cells across an entire dorsal to ventral region of the mouse retina. By analyzing cell fates in the context of their position in the tissue, we found that cones could be classified into two subclasses with a graded gene expression profile changing nonlinearly. This study exemplifies the benefits of quantitatively analyzing populations of cells in a tissue when classifying fate decisions.

In the mouse retina, we defined two cone subtypes, S-only cones and CEC cones, based strictly on opsin expression profiles. Interestingly, the population of S-only cones in the dorsal region have higher S-opsin expression than most S-only cones in the ventral region (**Fig. 5J**). These highly expressing S-only cones are found at a steady density across the D-V axis of the retina. It is possible that this subset of evenly distributed, high S-opsin expressing cells could comprise the “genuine” S-cones that connect to blue-cone bipolar neurons (Haverkamp et al., 2005). We were not able to mathematically distinguish genuine S-cones from the total population of S-only cones. Together, opsin expression and connectivity suggest three possible distinct cone subtypes: 1. CEC cones that do not connect to blue-cone bipolars, 2. “genuine” S-only cones that connect to blue-cone bipolars, and 3. S-only cones that do not connect to blue-cone bipolars. Examination of connectivity in conditions that perturb thyroid hormone signaling may inform the relationship between opsin expression and connectivity and their relationship to cone subtype.

We developed a mathematical model that described both the binary fate specification process of cones and the graded expression of opsins, all driven by an external gradient. Stochastic modeling of this complex process generated probability distributions that we used to compare with experimentally observed cell distributions to test hypotheses about the connections between cell fates. Stochastic modeling is now sufficiently mature to perform detailed simulations of tissue-level cell-fate decisions. Combining stochastic models with high-throughput microscopy is a powerful tool for helping to understand complex relationships in tissues.

These methods advance our understanding of how regulatory inputs influence complex cellular decisions to specify binary and graded cell fates within the same cell type in the same tissue. A next step will be to integrate more signaling inputs into our model for retinal development. Numerous signaling molecules are expressed in D-V gradients and are involved in retinal and cone cell development. Specifically, retinoic acid (RA) is an important morphogen that is expressed at high levels ventrally during development, and then at moderate levels in the dorsal region in the adult mouse (McCaffery et al., 1992; McCaffrery et al., 1993). Moreover, further studies would include integrating transcription factor binding partners of Thrβ2 into the model, as Thrβ2 acts as a homodimer and as a heterodimer with RXRγ (Roberts et al., 2006). This work represents an important first step towards modeling the complex network of interactions that guide binary and graded cell fate specification.

The retina provides an excellent paradigm to study how signaling inputs generate patterns in two dimensions. The next challenge will be developing models for patterning in more complex 3-dimensional neural tissue found in brain structures. Quantitative modeling has enormous potential to integrate multiple signals across a tissue and build networks to better understand and predict the outcomes of development when variables are changed, for instance in disease states.

## Materials and Methods

### Animals

Mice (strain C57BL/6) were housed under a 12 h light:12 h dark (T24) cycle at a temperature of 22°C with food and water *ad libitum*. Male and female mice at 2–8 months of age were housed in plastic translucent cages with steel-lined lids in an open room. Ambient room temperature and humidity were monitored daily and tightly controlled. Wild-type mice (C57BL/6; Jackson Laboratory), and *Thrb^tm2Df^* mutant mice (gift from the Forrest Lab) were used in this study. *Thrb^tm2Df^* mutant mice specifically knock out expression of Thrβ2, and leave Thrβ1 intact as previously described (Ng et al., 2001). All animals were handled in accordance with guidelines of the Animal Care and Use Committees of Johns Hopkins University. All efforts were made to minimize the pain and the number of animals used.

### Immunohistochemistry

Retinas were dissected in PBS, then fixed in fresh 4% formaldehyde and 5% sucrose in PBS for 1 hour. The dorsal portion of the retina was marked with a cut. Tissue was rinsed 3X for 15 min in PBS. Retinas were incubated for 2 hours in blocking solution (0.2-0.3% Trition X-100, 2-4% donkey serum in PBS). Retinas were incubated with primary antibodies in blocking solution overnight at 4°C. Retinas were washed 3X for 30 min in PBS, and then incubated with secondary antibodies in blocking solution for 2 hours at room temperature. At the end of staining, retinas were cut to lay flat on a slide, and were mounted for imaging in slow fade (S36940, Thermo Fisher Scientific).

### Antibodies

Primary antibodies were used at the following dilutions: goat anti-SW-opsin (1:200) (Santa Cruz Biotechnology), rabbit anti-LW/MW-opsins (1:200) (Millipore). All secondary antibodies were Alexa Fluor-conjugated (1:400) and made in donkey (Molecular Probes).

### Microscopy and image processing

Fluorescent images were acquired with a Zeiss LSM780 or LSM800 laser scanning confocal microscope. Confocal microscopy was performed with similar settings for laser power, photomultiplier gain and offset, and pinhole diameter. Whole retinas were imaged with a 10X objective, and maximum intensity projections of z-stacks (5–80 optical sections, 4.9 μm step size) were rendered to display all cones imaged in a single retina. Retinal strips were imaged with a 20X objective, and maximum intensity projections of z-stacks (5–80 optical sections, 1.10 μm step size) were rendered to display all cones imaged in a single retina. ΔThrβ2 retinas and wildtype retinas were taken at the same time and stained with the same batch of antibody, then imaged with the same laser intensity for comparison of opsin levels.

### Segmentation of cone cells from microscopy images

Microscopy images were analyzed using a custom parallel image processing pipeline in Biospark (Klein et al., 2017). Briefly, each fluorescence channel was first normalized and filtered to remove small bright features. Then, each remaining peak in fluorescence intensity was identified and an independent active contour segmentation (Marquez-Neila et al., 2014) was performed starting from the peak. If the resulting contour passed validation checks it was included in the list of segmented cone cells for the channel. Finally, cell boundaries were reconciled across both channels to obtain a complete list of identified cone cells. Full details are given in the SI Methods.

### Modeling cone cells fate decisions and opsin expression

Multiscale modeling of the retina strip was performed using a hybrid deterministic-stochastic method. Diffusion of T3 in the microenvironment of the retinal strip was modeled using the diffusion partial differential equation (PDE). The PDE was solved using an explicit finite difference method. The cone cells were modeled using the chemical master equation (CME) to describe the stochastic reaction scheme implementing the cell fate decision-making. The CME for each of the 23,760 cone cells was independently sampled using Gillespie’s stochastic simulation algorithm (Gillespie, 1977). Reconciliation between the CME trajectories and the PDE microenvironment was done using a time-stepping approach. Complete mathematical details of the model and simulation methods are available in the SI Methods. All simulations were performed using a custom solver added to the LMES software (Roberts et al., 2013), which is available on our website: https://www.robertslabjhu.info/home/software/lmes/.

## Data Availability

Immunofluorescence images (https://osf.io/e5ckg/) and analysis code (https://osf.io/b438a) are available in the Open Science Framework database.

## Supplementary Material

### Supplementary Text

#### 1 Methods

##### 1.1 Analysis of retina microscopy images

###### 1.1.1 Segmentation of individual photoreceptor cells in 200X images

To identify and segment individual cone cells in the immunostained fluorescence images, we developed an image processing pipeline similar to that used previously [1]. The analysis begins finding potential cone cells by looking for connected regions in our 200X fluorescence images. Because some cells are present in only the blue (S-opsin) or green (M-opsin) channels and some cells are present in both, we combine all potential cells into a joint list for additional analysis. Algorithm 1 outlines this first step of our processing pipeline.

###### Algorithm 1

Extract the subimage surrounding each potential cone cell.

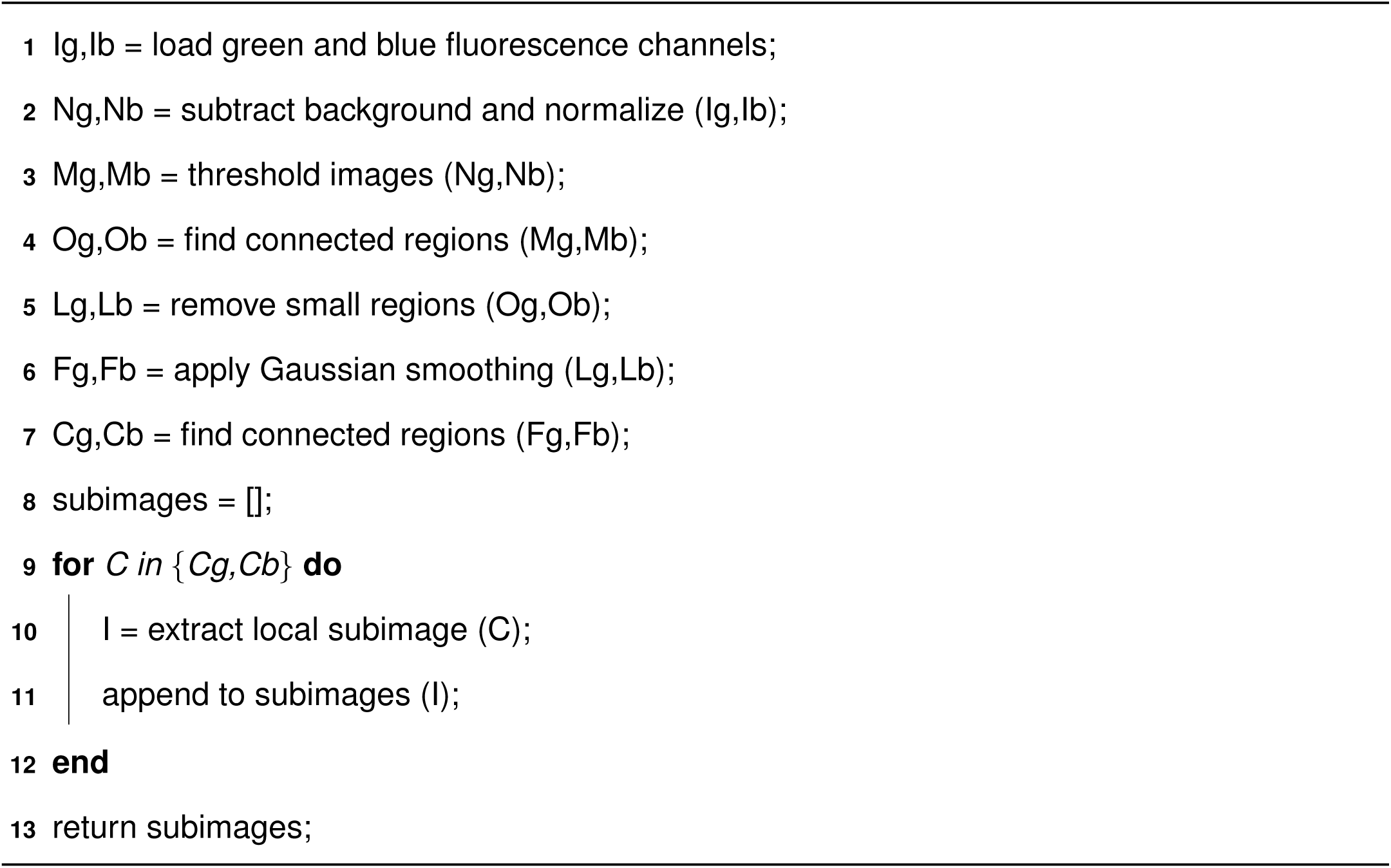

Next, we find the outline of each potential cell using an active contouring method known as morphological snakes [2]. Because we are segmenting tens-of-thousands of individual potential cells from each image, we perform this step of the pipeline in parallel using the Biospark framework [3], which is a data intensive parallel analysis package for Python. We perform validation of the segmented boundaries before classifying the object as a cone cell.

###### Algorithm 2

Identify and segment any cone cells in each subimage.

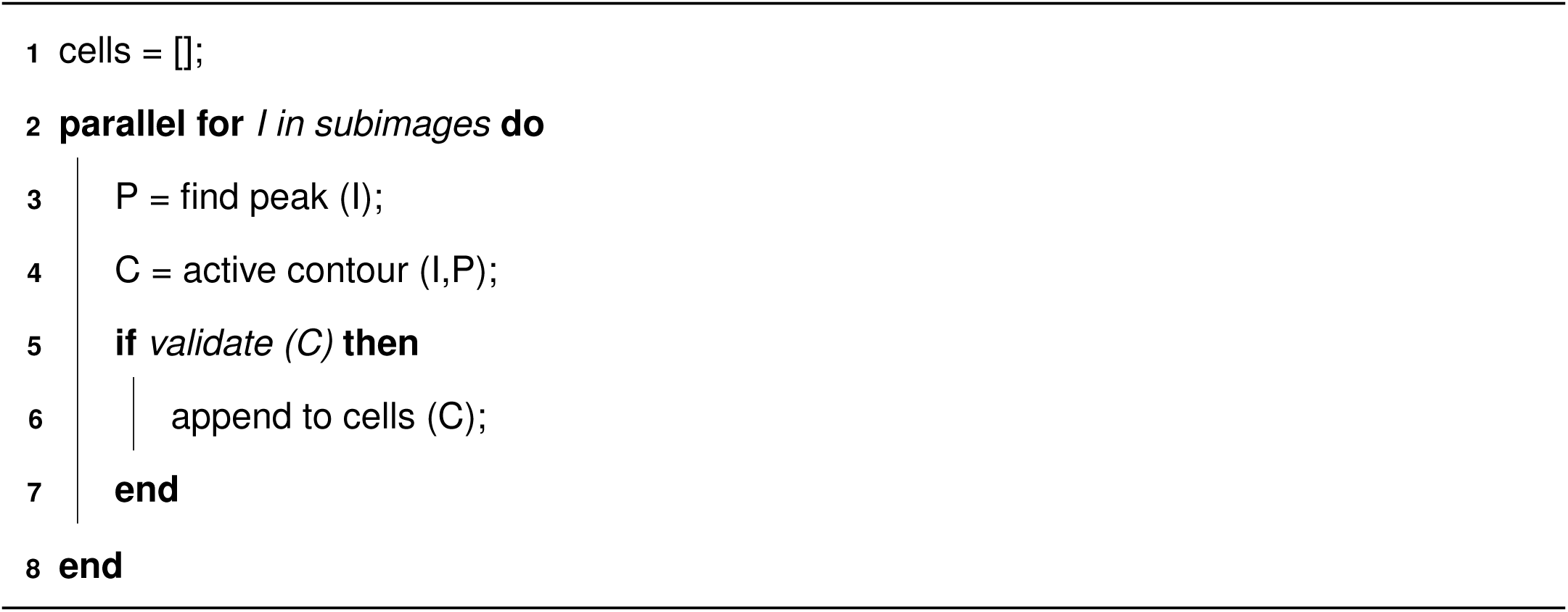

Finally, we perform a reconciliation step in which overlapping cells are merged and/or split to obtain an estimated final segmentation for the image. The pixel indices associated with each cell are stored so that the properties of each cone cell can be later calculated. All of the scripts and Jupyter notebooks implementing our analysis pipeline are available for download from our website: https://www.robertslabjhu.info/home/software/mouse_eye.

###### 1.1.2 Validation of segmentation results

We validated that our segmentation algorithm produced results similar to human annotators by comparing manually and automatically generated statistics from representative samples of our data set. We picked seven different regions and manually counted the number of S-opsin only, M-opsin only, and coexpressing cells. We then analyzed the same regions using our segmentation algorithm and obtained the automatically generated classifications.

Table S2 shows the counts of cell types from these regions from both human and computer annotations. Overall, there is excellent agreement in the relative abundance of the different cell types. The absolute counts have some systematic difference, with the automatic segmentation typically identifying more cells than human annotators. This is mostly due to what appear to be single long cells that are split into multiple bright pieces separated by low fluorescence breaks. Human annotators tend to regard the trace as a single long cell, while the automatic segmentation tends to identify multiple smaller cells. Importantly, we do not know the true underlying cell morphology, so we cannot generally say whether the human or automatic annotators are more accurate. In any case, since the cell parts are identified correctly and the segmentation is consistent across retinas, we expect these minor difference to have no impact on our results.

##### 1.2 Modeling of photoreceptor cells

###### 1.2.1 Modeling cone cell fate determination and opsin expression in a retinal strip

To model the cone fate decisions in a large retinal strip we use a combination of stochastic and deterministic modeling. We start with a three-dimensional volume 5 mm long in the X dimension, 1 mm wide in the Y dimension, and 5 *µ*m in the Z dimension representing the microenvironment of the dorsal-ventral (DV) strip. The small z dimension make this an effectively two-dimensional system and we include *z* below only for completeness. Within this volume we model diffusion of thyroid hormone (T3) using the deterministic diffusion equation:

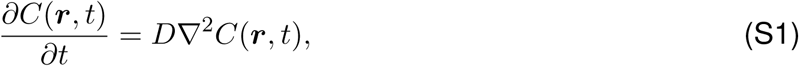

where *C*(***r****, t*) is the concentration of T3 at position ***r*** and time *t*, *D* is the diffusion coefficient used for T3, and *∇*^2^ is the Laplace operator 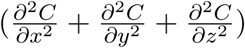. See Table S3 for all parameter values used.

We numerically solve the diffusion partial differential equation (PDE) using a explicit finite difference method with grid spacing *dx* and a time step *dt* = (*dx*^2^)*/*(2 *·* 6*D*), where the extra factor of 2 in the denominator ensures numerical stability. We fix the concentration at the *X* = 0 boundary to *C_hi_* and at the opposite boundary to *C_lo_* to establish a stationary concentration gradient in the X dimension. The Y and Z boundaries are taken to be reflective. We initialize the concentrations *C*(***r***, 0) according to a linear decrease in ***r*** to follow to the boundary conditions.

Within the microenvironment, we place 23,760 individual photoreceptor cells spaced on a hexagonal grid spanning the X-Y plane, with center-to-center distance *d_cell−cell_* and with a radius *r_cell_*. Each cell is modeled independently using the chemical master equation (CME) describing the probability to have a particular count of each species:

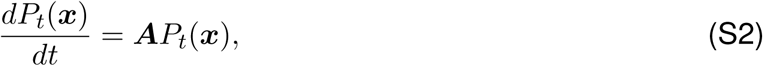

where *P_t_*(***x***) is the probability for a cell to have a particular state vector ***x*** giving the count for each chemical species and ***A*** is a transition matrix describing all of the reactions between the chemical species.

Within each photoreceptor cell a series of reactions describing the fate of the cell and also opsin expression take place. First, cells contain thyroid hormone receptors THR*β*2. T3 can bind reversibly to THR*β*2 to switch it to an activated state THR*β*2*^∗^*:

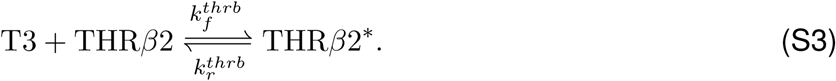

Note that T3 also diffusing across the PDE microenvironment and the value of T3 is synchronized between the PDE and CME models. The synchronization procedure is discussed below.

Next, the cells switch between three cell fates. FD(U) cells are undifferentiated and do not express opsins, FD(S) cells occupy an S-only cone cell fate with high expression of only S-opsin, and FD(C) cells are typical cone cells that express some combination of S- and M-opsin depending on various factors. The transition between cone cell fates in real cells depends on a number of unknown fate determining steps. Little is known about these steps, but the fate decisions appear to be stable. Therefore, we model photoreceptor fate decision-making as barrier crossing process with a variable number of cooperative steps *n*.

The FD(U) *↔* FD(S) transition is described by

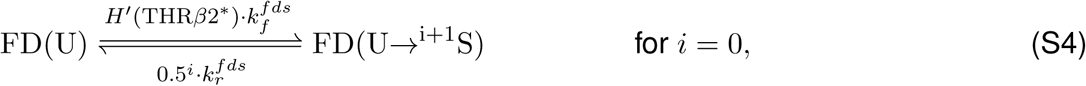

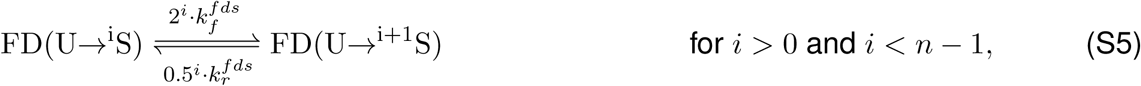

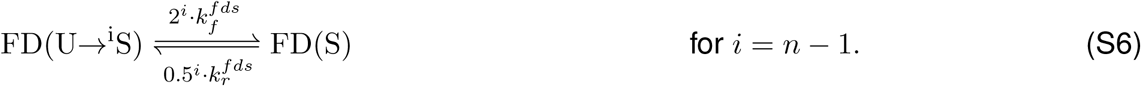

Here, the syntax FD(U*→^i^*S) denotes a cell that has progressed *i* steps along the path from the FD(U) to FD(S) fate. Also, *H^t^* is an inhibiting Hill-like kinetic function defined by

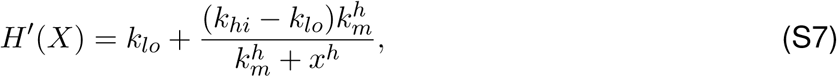

with *k_lo_* and *k_hi_* the lower and upper limits of the kinetic process, respectively, *k_m_* the midpoint of the transition, and *h* the Hill exponent giving the cooperativity of the transition. Likewise, the FD(U) *↔* FD(C) transition is described by

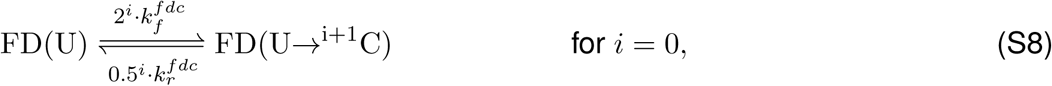

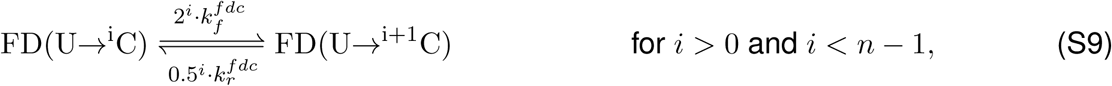

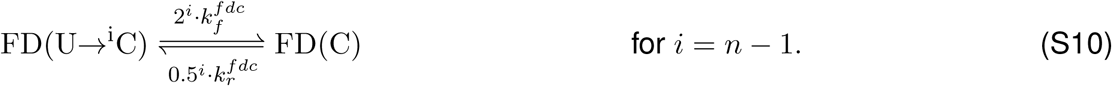

Then, both S-opsin and M-opsin proteins can be expressed by photoreceptor cells, depending on the cell type and the local concentration of T3. We model opsin expression using the following kinetic equations

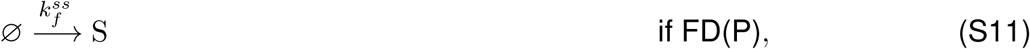

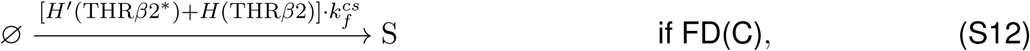

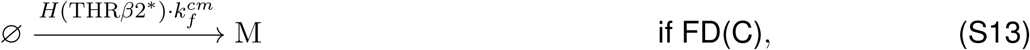

*H* is an activating Hill-like kinetic function

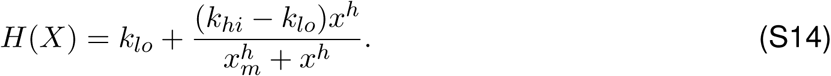

Finally, both opsins can be degraded according to

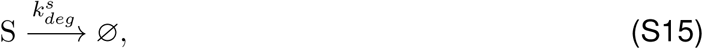

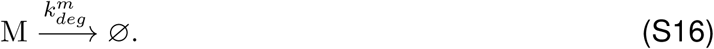

We initialize each cell at *t* = 0 to the FD(U) state with a copy number of THR*β*2 proteins independently sampled from a Gaussian distribution with a mean concentration of *µ_thrb_* and a variance of 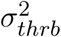. The number of T3 molecules available to the cell is initialized according to the T3 concentration at the cell’s D-V position. Likewise, the fraction of activated THR*β*2 is initialized to its equilibrium value according to the cell’s D-V position. All opsin counts are initialized to zero. We then model the stochastic time evolution of each cell using the standard Gillespie stochastic simulation algorithm (SSA) [4, 5].

###### 1.2.2 Microenvironment modeling of combined PDE and CME dynamics

Since we are performing two parallel simulations, PDE and CME, we need to partition the molecules between them. Each cell has a volume smaller than a PDE subvolume and each cell is assumed to be completely contained within a single subvolume. We initialize the T3 molecule count in the extracellular space of each cell to be the rounded number of molecules corresponding to the cell’s extracellular volume multiplied by the PDE subvolume’s concentration. In this way T3 molecules are represented in each simulation.

To integrate the PDE and CME dynamics, we implemented a parallel time-stepping approach. We divide time into discrete synchronization intervals Δ*t* and evolve overall time according to

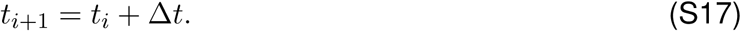

During each Δ*t* we update the state of each cell using the SSA and the concentration of each PDE subvolume using a finite difference algorithm. The loss or gain in the number of extracellular T3 molecules in the stochastic simulation during the time step is tracked. This gain or loss is converted into a concentration flux and applied to the PDE subvolume during the next time step. Simultaneously, the number of extracellular T3 molecules in the stochastic model is reset for the next time step according to the new subvolume concentration. Our method conserves mass and has good error characteristics as long as the T3 flux at each time step is of the order of a few molecules. The method is implemented as the “microenvironment” solver in our LMES software and is freely available on our website: https://www.robertslabjhu.info/home/software/lmes.

###### 1.2.3 Parameterization of retinal strip microenvironment model

We parameterized our model by globally fitting the model parameters to five different dorsal-ventral (D-V) data sets: (1) the fraction of all cells expressing M-opsin, (2) the fraction of all cells expressing S-opsin, (3) the fraction of FD(S) cells expressing only S-opsin, (4) the per cell M-opsin expression level, and (5) the per cell S-opsin expression level. Because our data were collected from multiple retinas, we first fit the raw data to functions that we could use to describe a hypothetical mean retina.

We describe the fraction of cells in the various subfates as (modified) Hill-like functions. The fraction of cells expressing S-opsin as a function of D-V position *x* is described by:

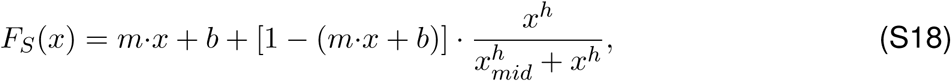

where *m* and *b* are the slope and x-intercept of a baseline fraction, respectively, *x_mid_* is the midpoint of the transition, and *h* is the Hill coefficient. Likewise, the fraction of M-opsin expressing cells is given by:

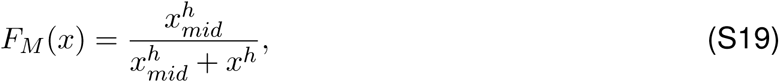

and the fraction of FD(S) cells is given by:

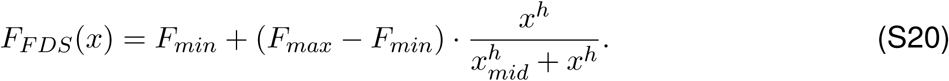

Figures S1+S3 show the fits of these functions to the raw data for the various retinas. We then took the mean of the various parameters to construct a hypothetical mean retina. Figure S13 shows the fraction of cells in these states as a function of D-V position in our mean retina.

We describe the mean per cell expression level of M- and S-opsin as piecewise linear functions with a low and a high limit separated by a biphasic region with two different slopes:

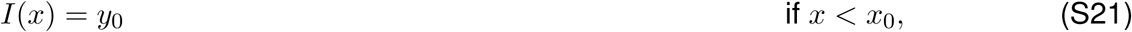

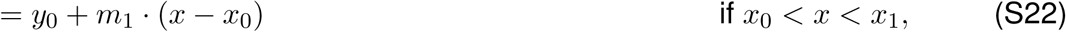

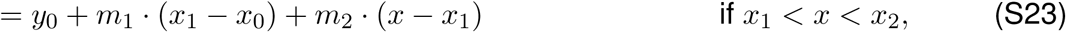

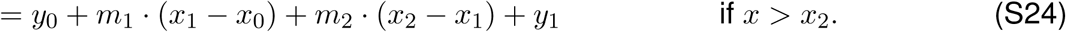

Here, *y*_0_ and *y*_1_ are the left and right baselines, *x*_0_, *x*_1_, and *x*_2_ are the D-V points where the slope changes, and *m*_1_ and *m*_2_ are the two slopes. Figure S7 shows the fit to the M- and S-opsin expression for the various experimental retinas. Figure S13 shows the values of these functions for our hypothetical mean retina.

Once we had a hypothetical mean retina, we used it to parameterize our model. We derived expressions for the mean value of the various observables as a function of D-V position by solving the deterministic system of ordinary differential equations described by Equations S3-S16. We then used non-linear least squares with Nelder-Mead minimization to globally optimize the parameters using all five data sets. Table S3 gives the best fits values for the free parameters and Figure S14 shows a comparison of the deterministic model calculated using the best-fit parameters against our hypothetical mean retina.

### Supplementary Tables

**Table S1:** Retina image names and genotypes.

| Retina | Filename | Genotype | Num. Cells |
| --- | --- | --- | --- |
| R07 | 170505_2M_WT_F1_Left_20x_Stitch_MIP | WT | 65467 |
| R08 | 170505_2M_WT_F1_Right_20x-Stitch-MIP | WT | 50437 |
| R09 | 170505_2M_WT_F2_Left_20x-Stitch-MIP | WT | 34381 |
| R22 | 170817_pregnantMALE_CONTROL_BL6_1_20x-Stitch-MIP_c1+2 | WT | 40660 |
| R25 | 171026_WT_F1_Left_20x-Stitch-MIP | WT | 29831 |
| R26 | 171026_WT_F1_Right_20x-Stitch-MIP | WT | 29962 |
| R31 | 180123_ThrB2_KO_4M_DV_Right_slide1_20x_Stitch-MIP | ΔTHRβ2 | 10781 |
| R32 | 180123_ThrB2_KO_4M_DV_Right_slide2_20x-Stitch-MIP | ΔTHRβ2 | 10508 |
| R33 | 180123_ThrB2_KO_4M_whole_Right_slide1_20X-Stitch-MIP | ΔTHRβ2 | 15570 |

**Table S2:** The number of cells identified per subtype by hand (H) and computer (C) annotation.

| Retina Section Filename | S- & M-opsin (H/C) | S-only (H/C) | M-only (H/C) |
| --- | --- | --- | --- |
| 171026_WT_F1_Left_20x-Stitch-MIP-DORSAL | 13/23 | 4/6 | 584/753 |
| 171026_WT_F2p_Left_780_20x-Stitch-MIP-VENTRAL | 267/313 | 82/70 | 2/1 |
| 171026_WT_F2p_Left_780_20x-Stitch-MIP-CENTER | 186/277 | 20/26 | 0/1 |
| 171026_WT_F2p_Left_780_20x-Stitch-MIP-DORSAL | 4/5 | 10/13 | 427/377 |
| 171026_WT_F2p_Right_800_20x-Stitch-MIP-VENTRAL | 380/783 | 198/303 | 0/0 |
| 171026_WT_F2p_Right_800_20x-Stitch-MIP-CENTER | 515/1060 | 64/81 | 8/1 |
| 171026_WT_F2p_Right_800_20x-Stitch-MIP-DORSAL | 17/29 | 3/4 | 259/264 |

**Table S3:**
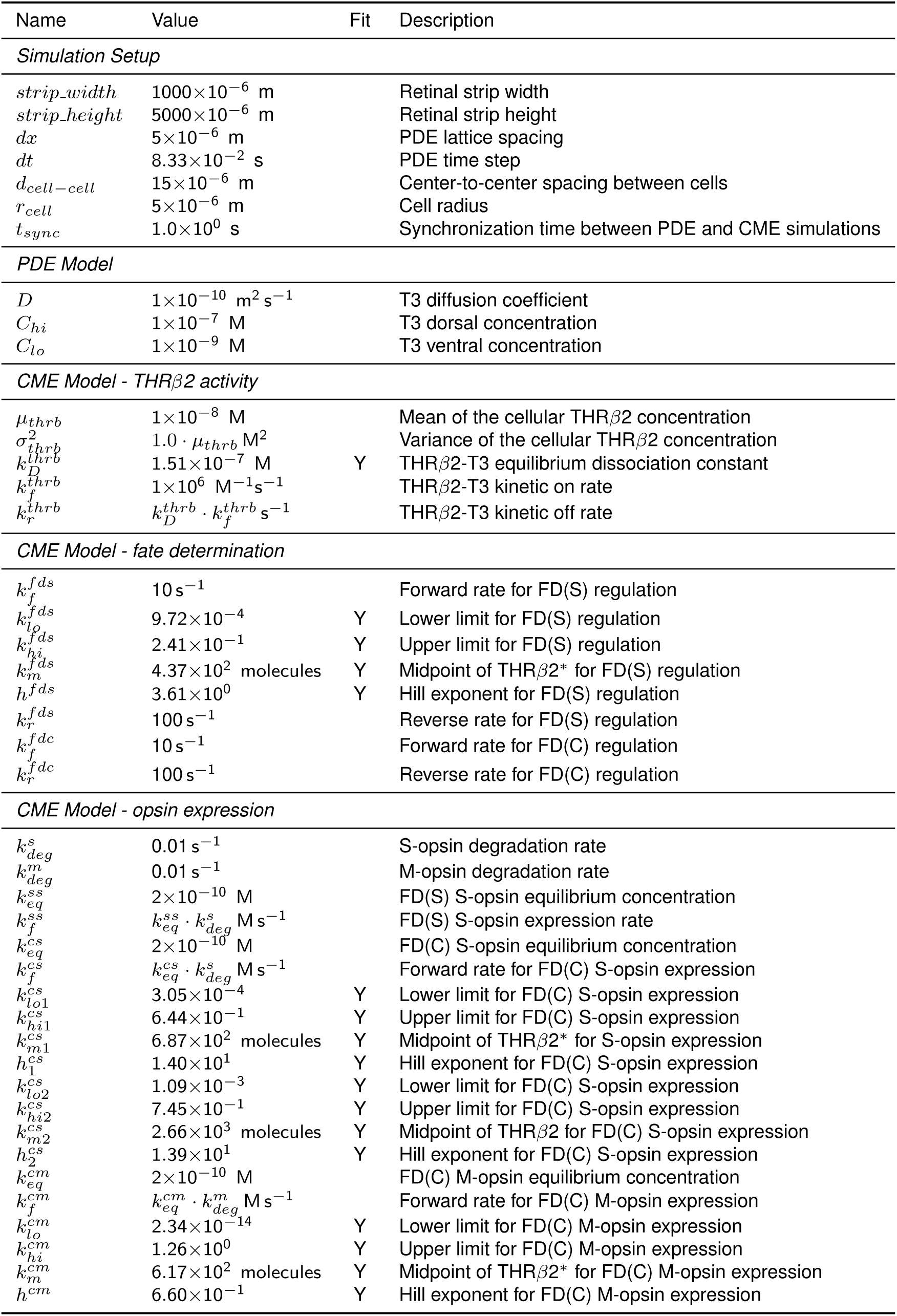
Parameters used in the microenvironment model.

### Supplementary Figures

**Figure S1:**
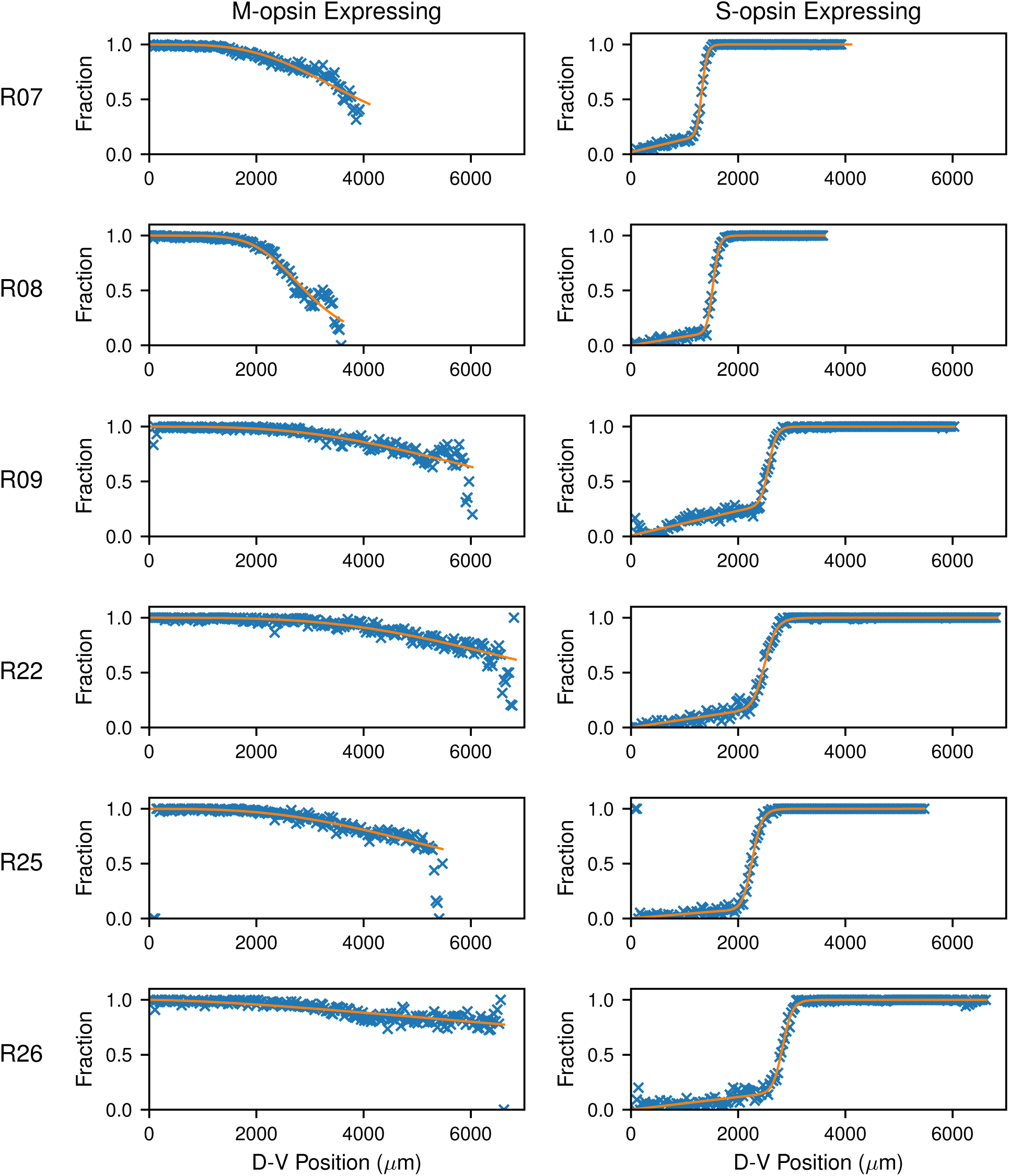
Fitting of cell expression data. Fraction of cells expressing (left) M-opsin and (right) S-opsin by position along the D-V axis. The data from the microscopy analysis (x) are overlaid with the best fit (line) to a fitting function (see text). Rows show different retinas (RXX).

**Figure S2:**
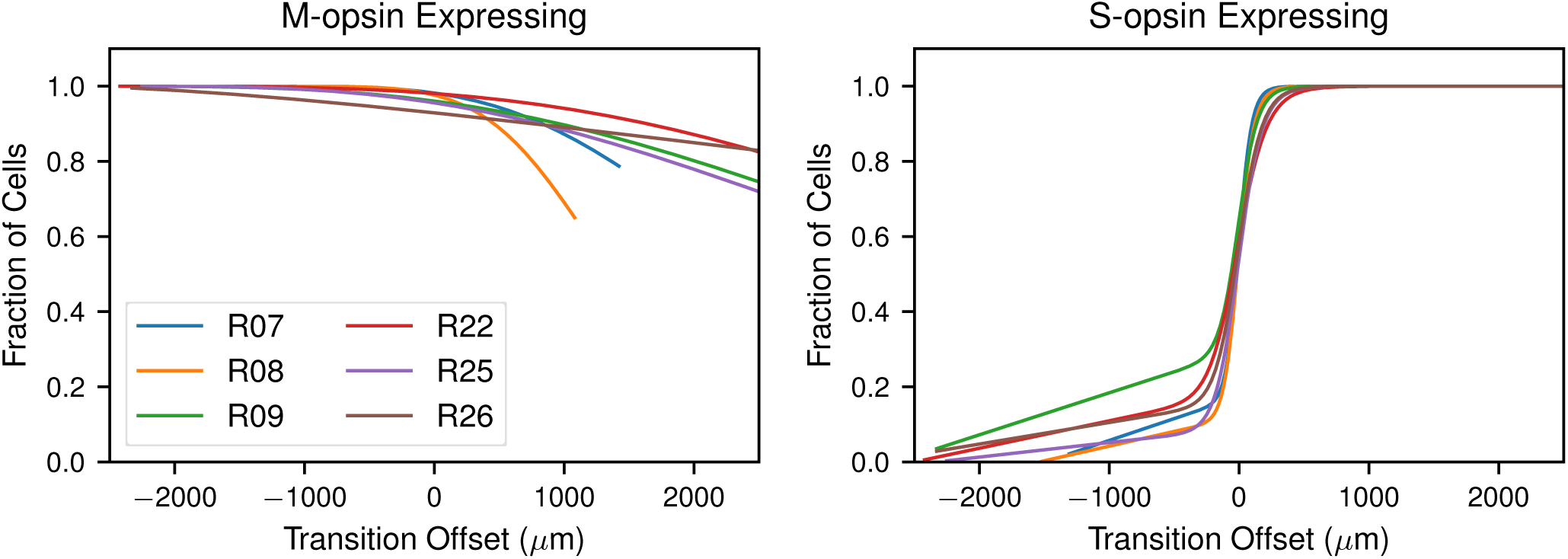
Comparison of D-V profiles between retinas. Overlap of the fraction of cells expressing (left) M-opsin and (right) S-opsin aligned to the transition midpoint as determined from the S-opsin expression profile.

**Figure S3:**
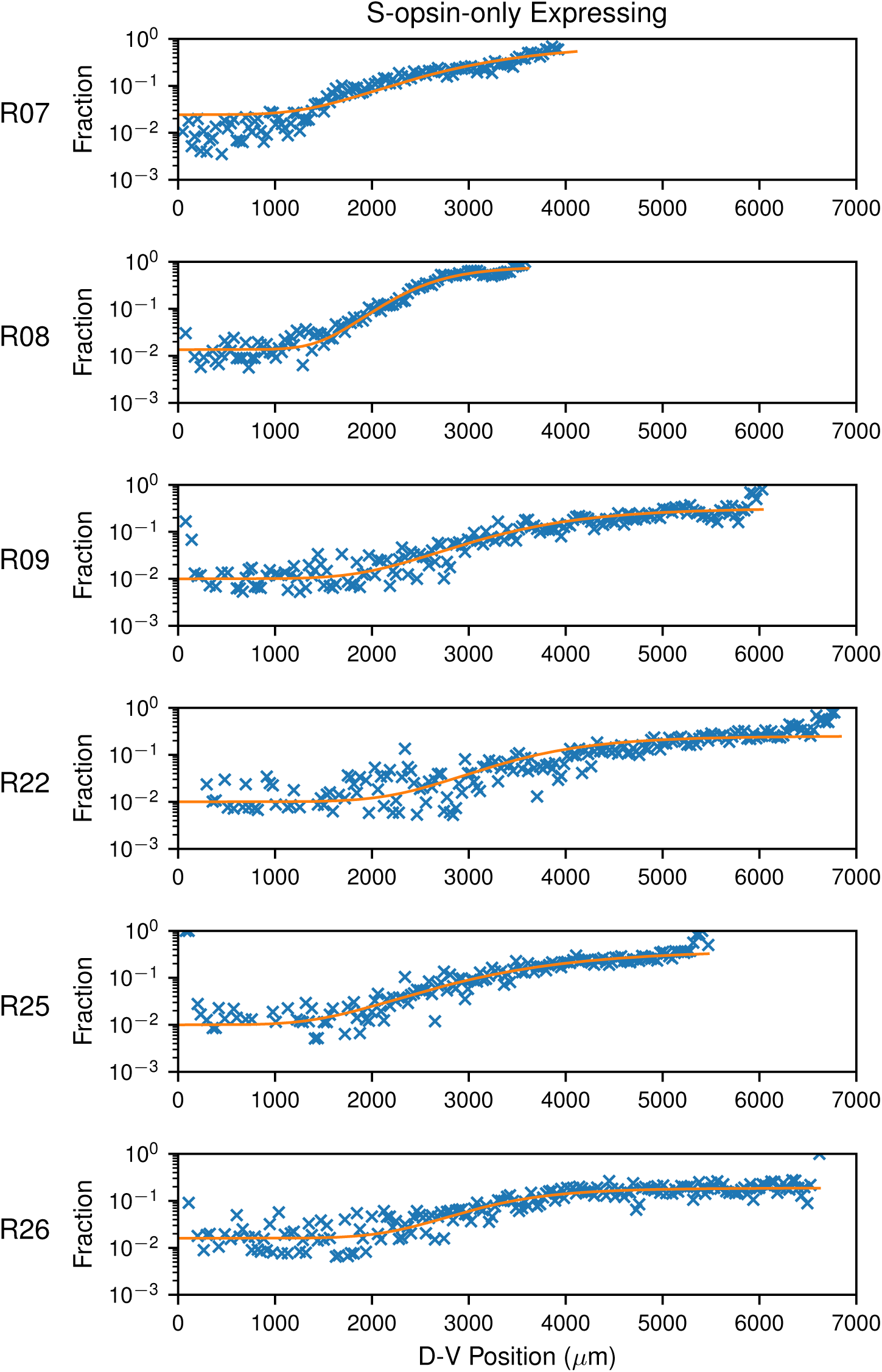
S-only cell fraction. Fraction of cells expressing only S-opsin by position along the D-V axis. The data from the microscopy analysis (x) are overlaid with the best fit (line) to a fitting function (see text). Rows show different retinas (RXX).

**Figure S4:**
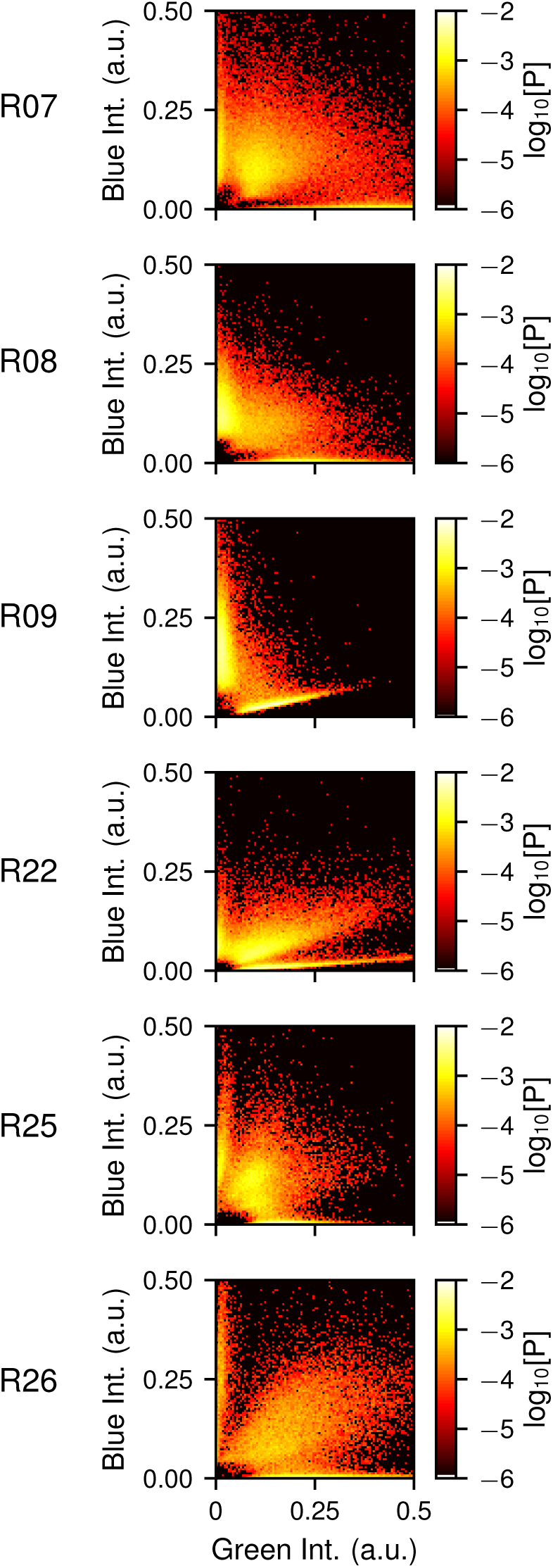
Correlation between S- and M-opsin in retinal cells. Joint probability distributions for the abundance of S-opsin (blue intensity) and M-opsin (green intensity) in cells. Rows show different retinas (RXX). Colors range from log_10_[P] = *−*2 (white/yellow) to log_10_[P] = *−*6 (red/black).

**Figure S5:**
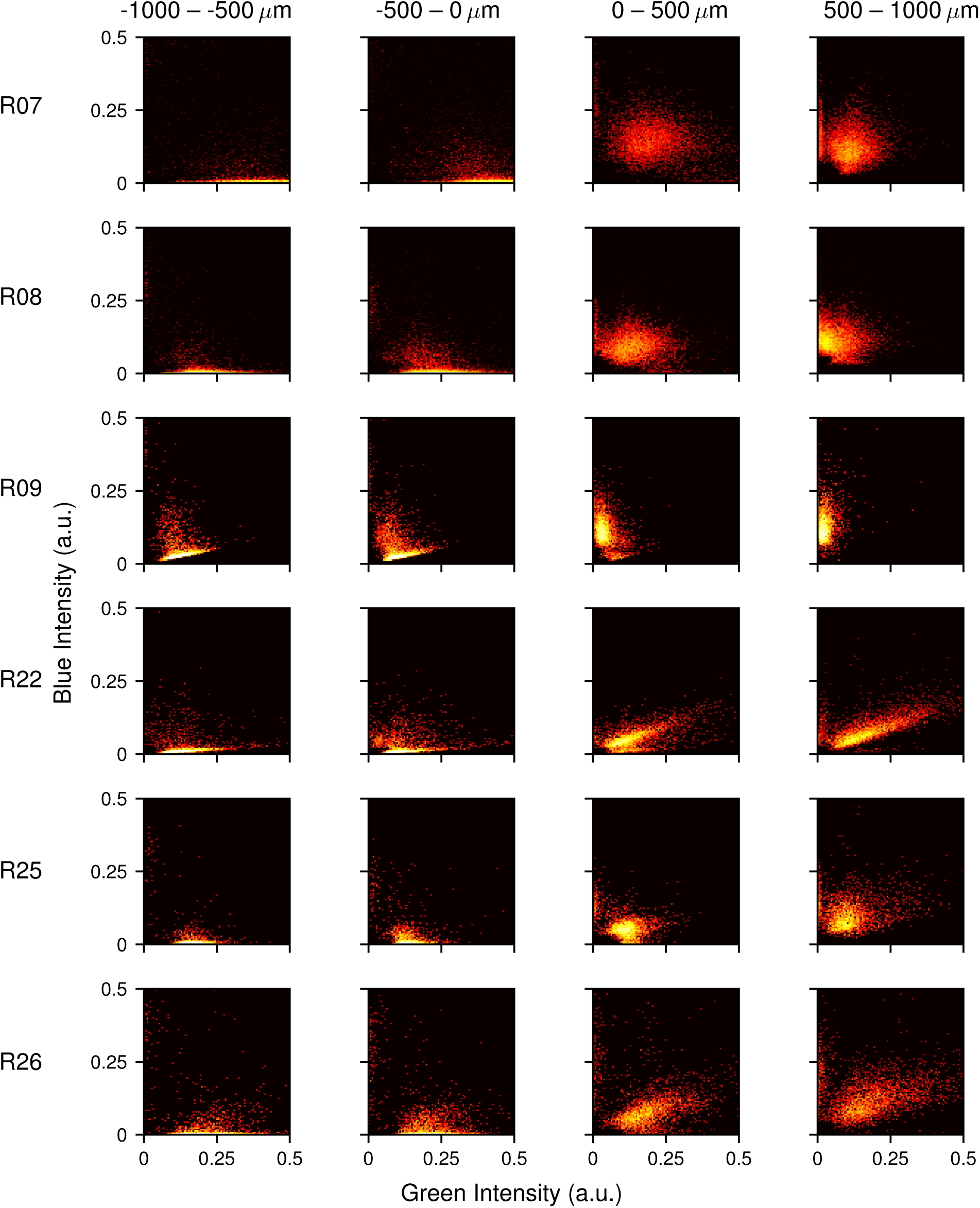
Correlation between S- and M-opsin in retinal cells. Joint probability distributions for the abundance of S-opsin (blue intensity) and M-opsin (green intensity) in cells. Columns show cells binned from four different regions according to distance from the transition midpoint. Rows show different retinas (RXX). Colors range from log_10_[P] = *−*2 (white/yellow) to log_10_[P] = *−*4 (red/black).

**Figure S6:**
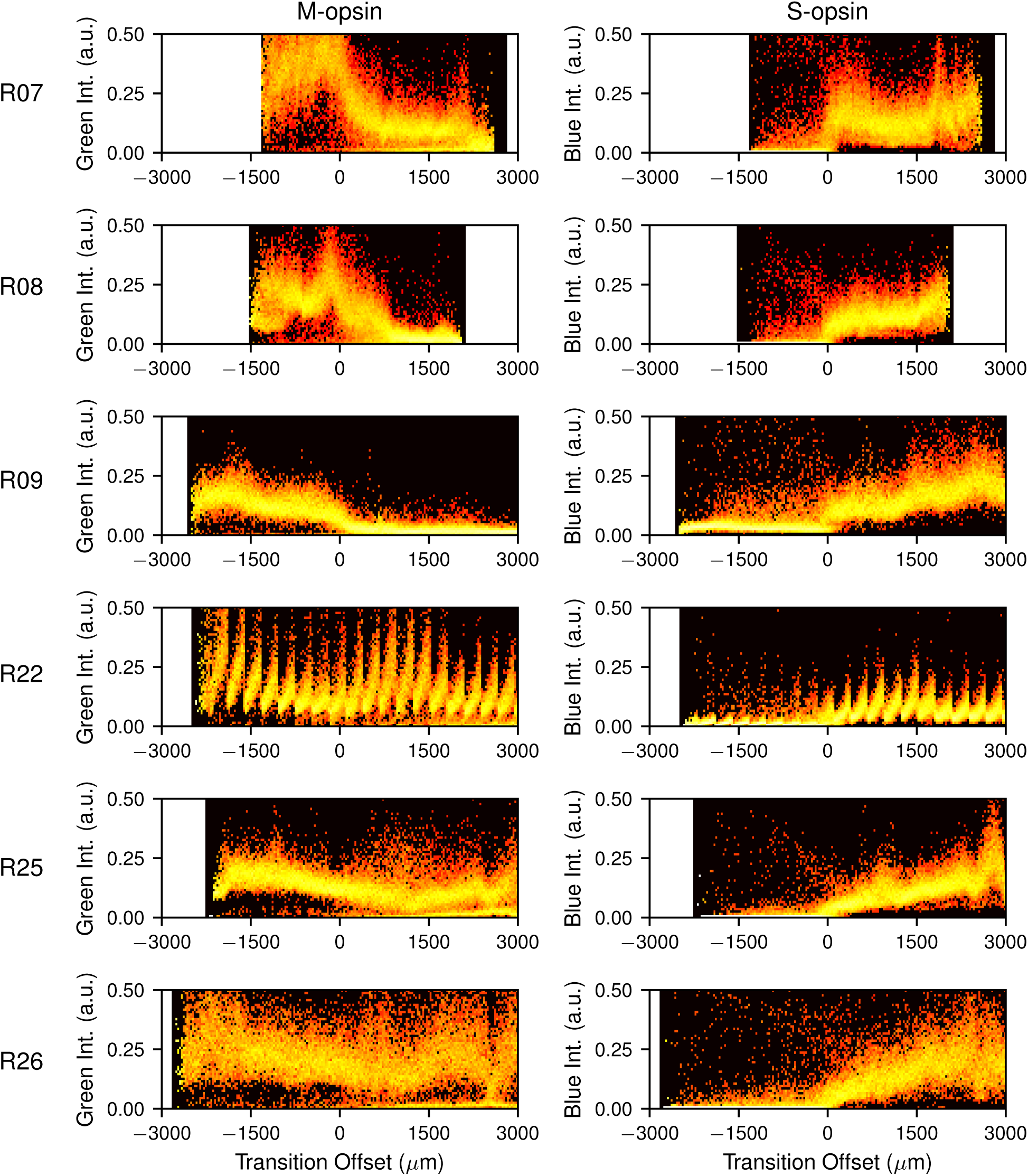
Expression of S- and M-opsin in retinal cells. Probability distribution for the abundance of (left) M-opsin and (right) S-opsin in cells by distance from the transition midpoint. Rows show different retinas (RXX). Colors range from log_10_[P] = 0 (white/yellow) to log_10_[P] = *−*4 (red/black).

**Figure S7:**
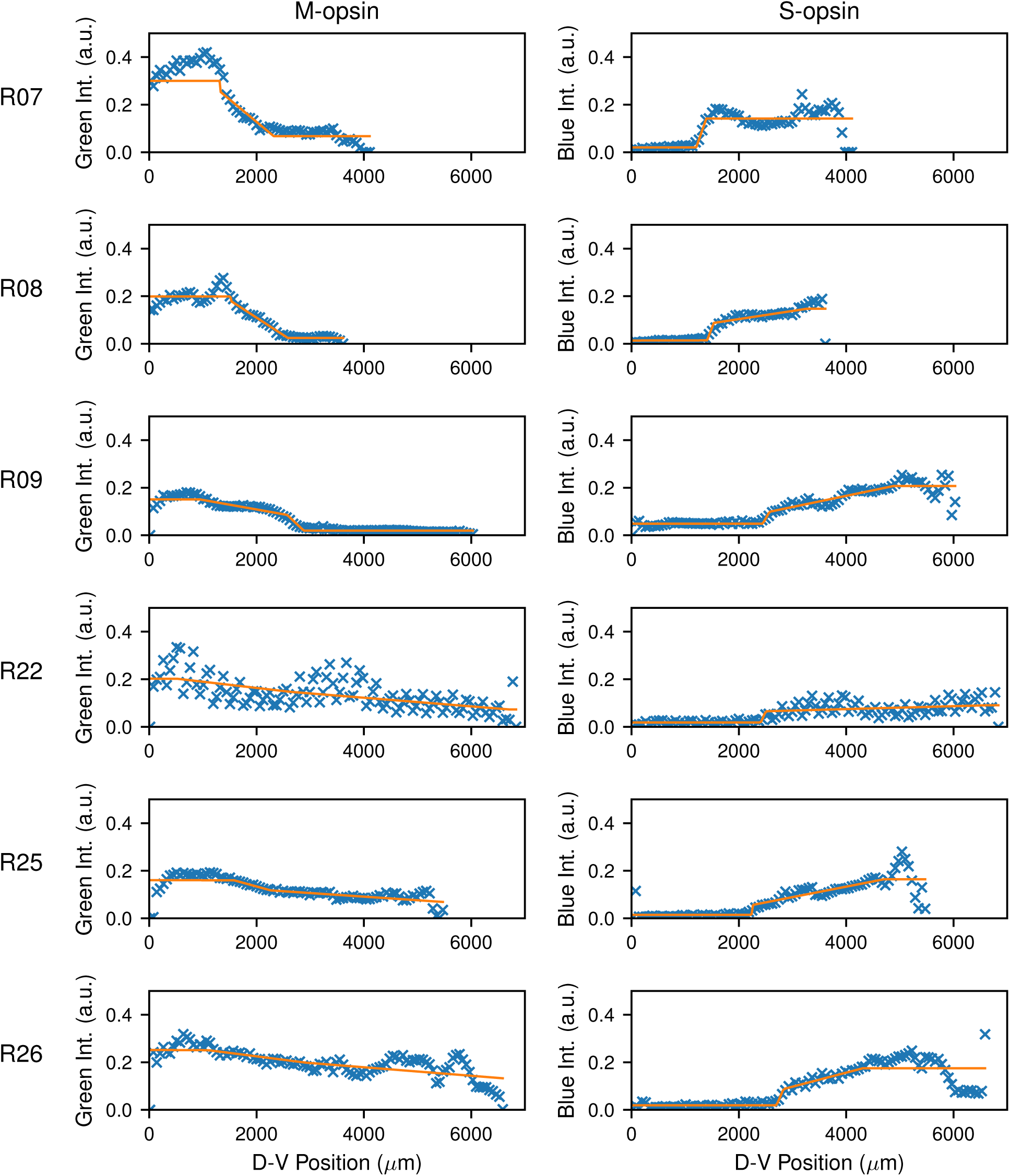
Fitting of cell expression intensity data. Mean intensity in all cells of (left) M-opsin and (right) S-opsin by position along the D-V axis. The data from the microscopy analysis (x) are overlaid with the best fit (line) to a fitting function (see text). Rows show different retinas (RXX).

**Figure S8:**
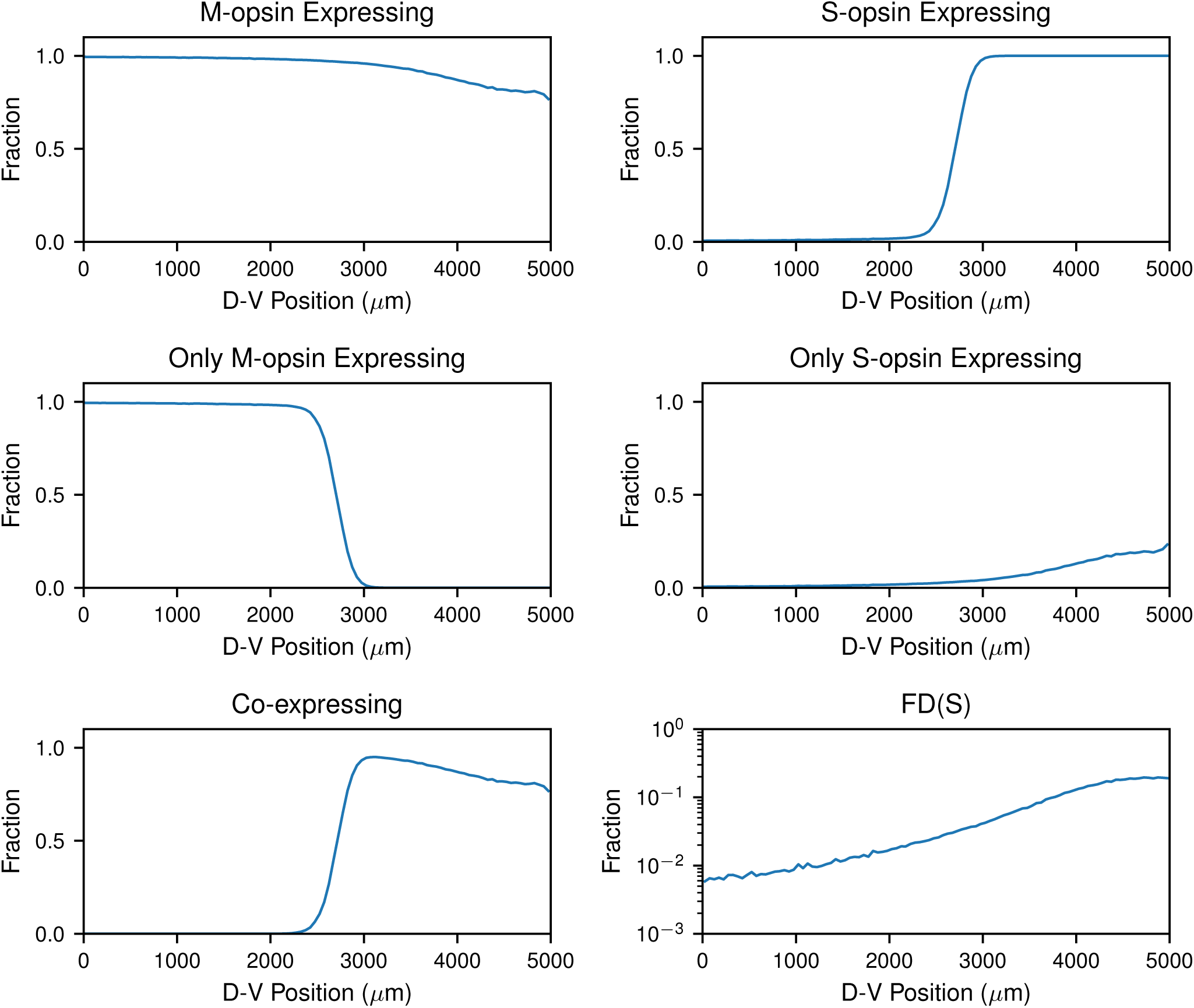
Expression in modeled cell populations. Mean fraction of cells in various cell populations along the D-V axis from numerical simulations of the model. Plots show the mean value computed from 100 independent simulations.

**Figure S9:**
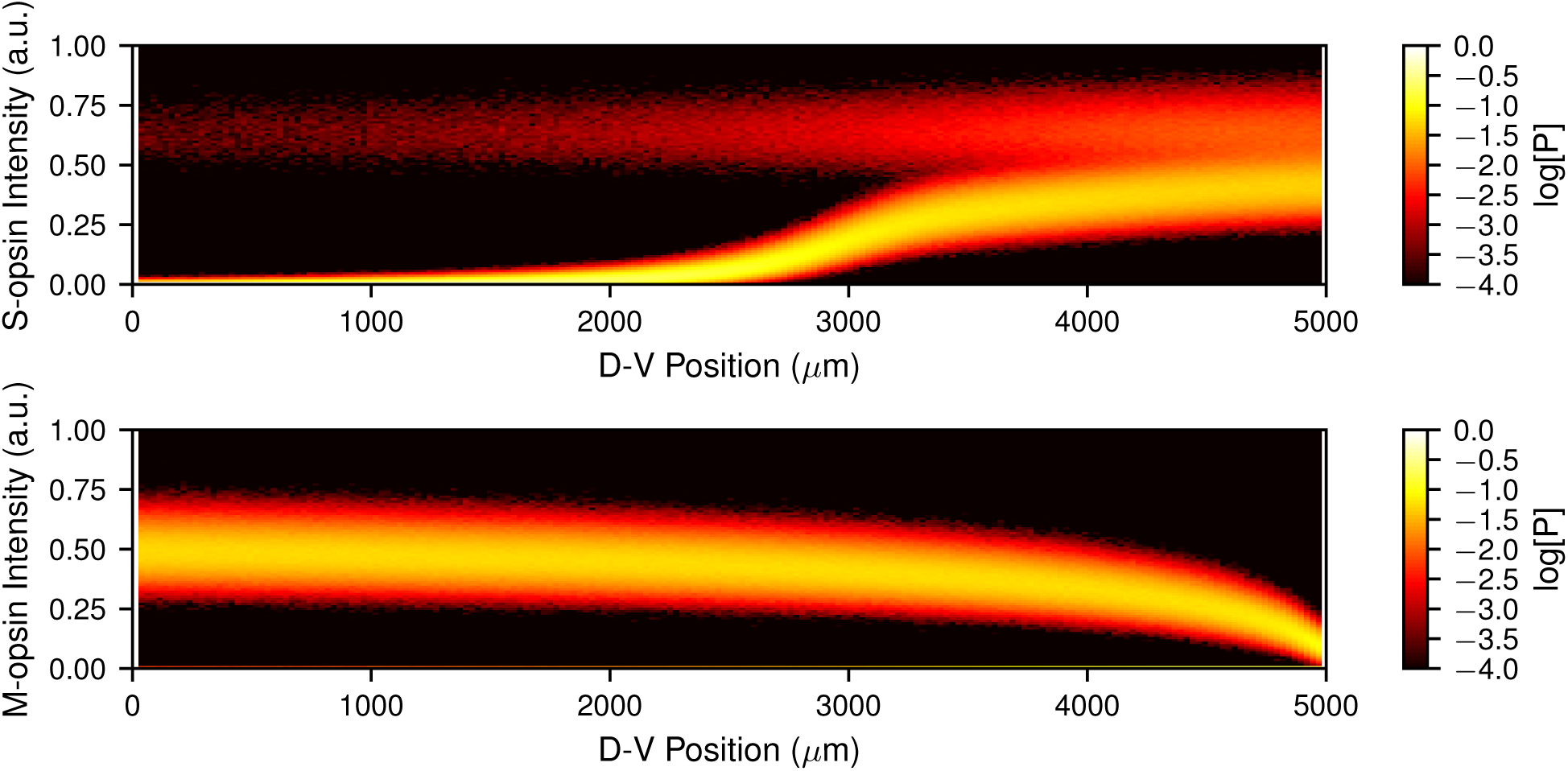
Opsin concentrations in modeled cells. Probability distribution of the abundance of S-opsin (blue intensity) and M-opsin (green intensity) in cells along the D-V axis from numerical simulations of the model. Distributions were computed from 100 independent simulations.

**Figure S10:**
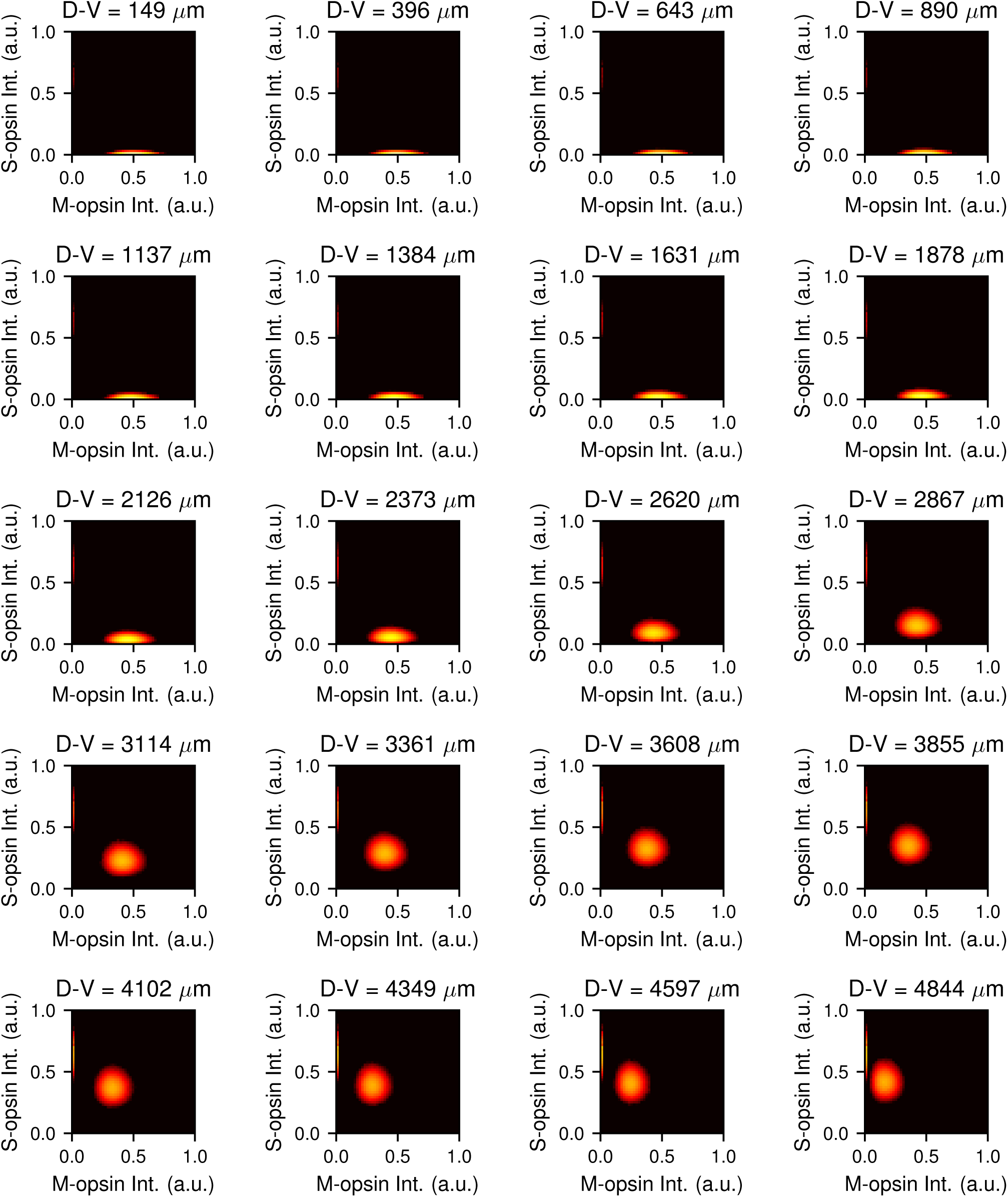
Correlation between S- and M-opsin in modeled cells. Joint probability distributions for the abundance of S-opsin (blue intensity) and M-opsin (green intensity) in cells located in ~250*µ*m wide bins along the D-V axis. Colors range from log_10_[P] = −2 (white/yellow) to log_10_[P] = −5 (red/black). Distributions were computed from 100 independent simulations. The low density tails leading to 0,0 are from cells that were sampled during the process of switching phenotypes.

**Figure S11:**
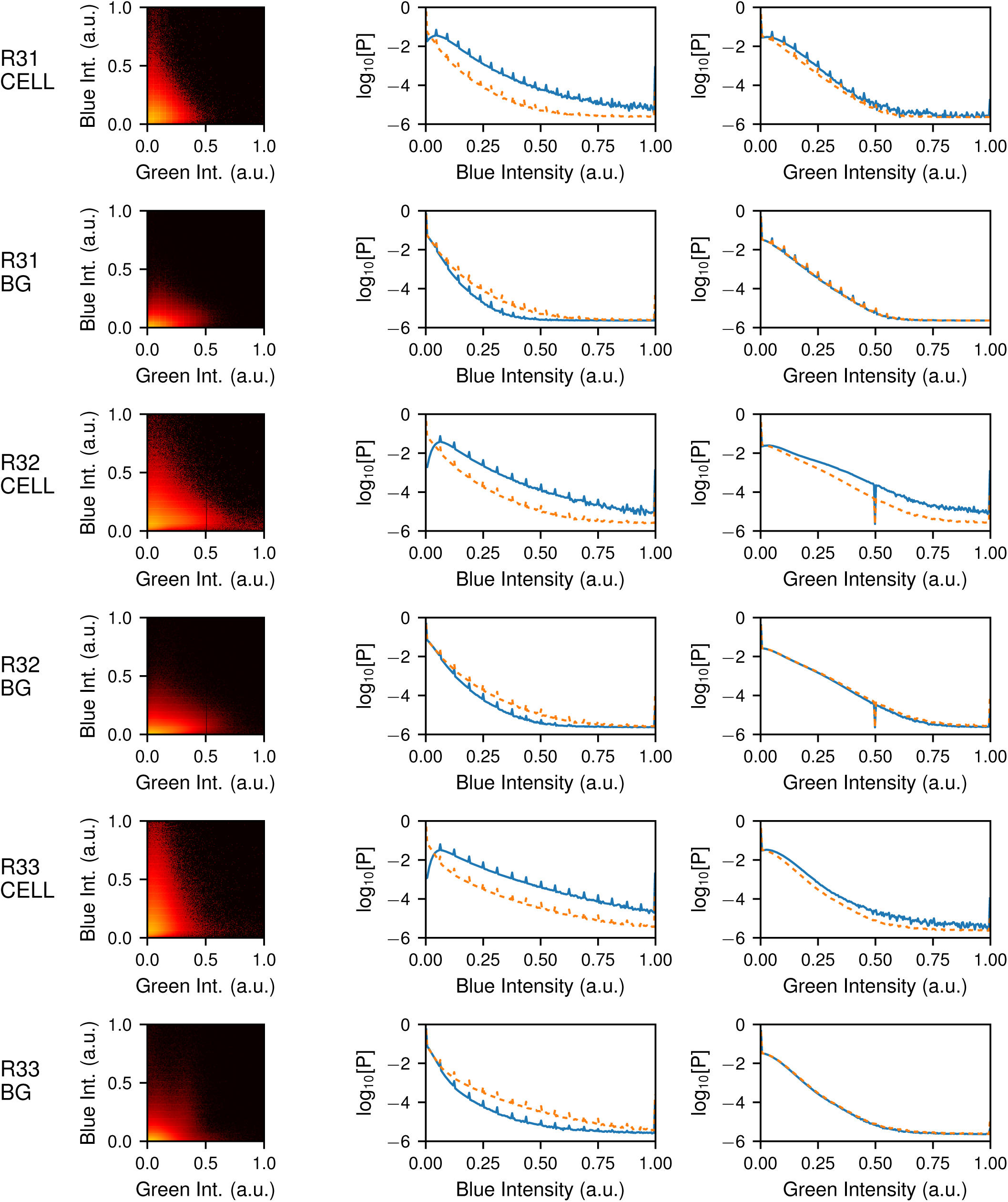
Analysis of pixel intensities in images of ΔTHR*β*2 cells. (left) Joint probability distribution of the blue and green intensity of pixels located either inside of cell boundaries (RXX CELL) or the background outside of cells (RXX BG) as indicated. Colors range from log_10_[P] = 0 (white/yellow) to log_10_[P] = −8 (red/black). (center) Probability for a pixel of the indicated type to have a particular blue intensity (solid line) compared with the distribution for all pixels (dashed line). (right) The same for green intensity. ΔTHR*β*2 cells do not exhibit green expression above background.

**Figure S12:**
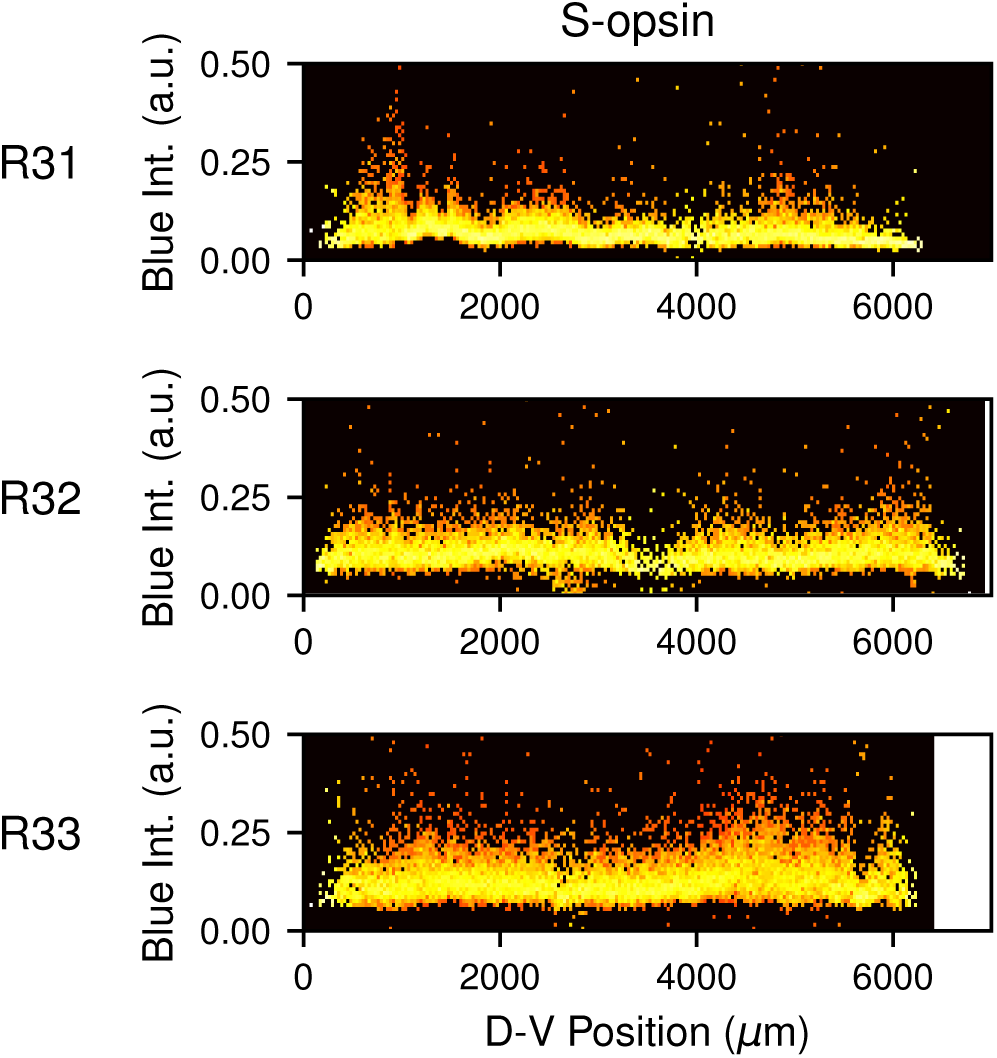
Expression of S-opsin in ΔTHR*β*2 retinal cells. Probability distribution for the abundance of S-opsin in cells by distance along the D-V axis. Rows show different ΔTHR*β*2 retinas (RXX). Colors range from log_10_[P] = 0 (white/yellow) to log_10_[P] = *−*4 (red/black).

**Figure S13:**
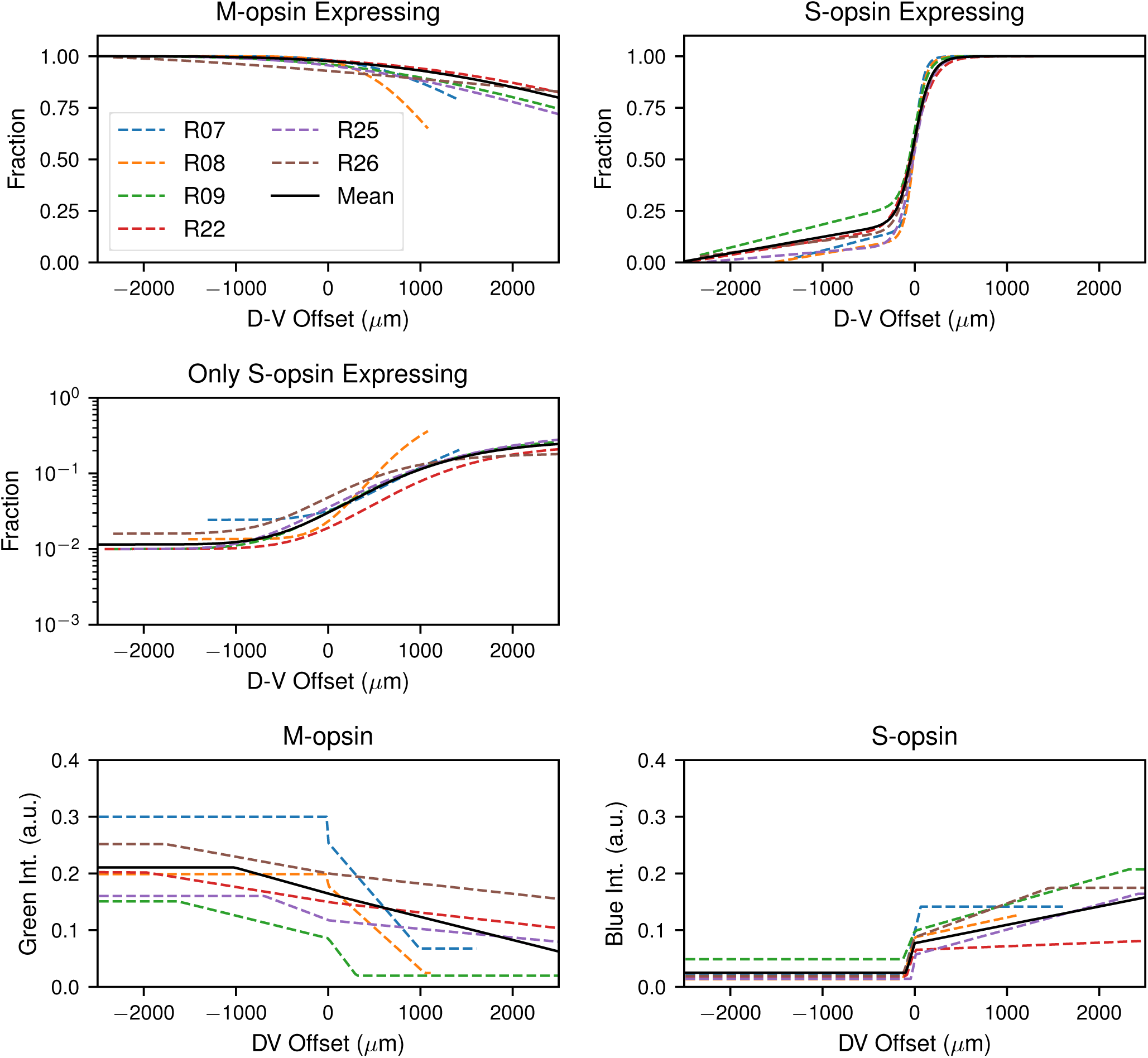
Mean retina description. Comparison of the fits for individual retinas (dashed lines) with our hypothetical mean retina used for model parameterization (solid line) along the D-V axis. The top row shows a comparison of the fraction of cells expressing M- and S-opsin, respectively. The middle row shows the fraction of FD(S) cells. The bottom row shows the mean M- and S-opsin expression intensity, respectively.

**Figure S14:**
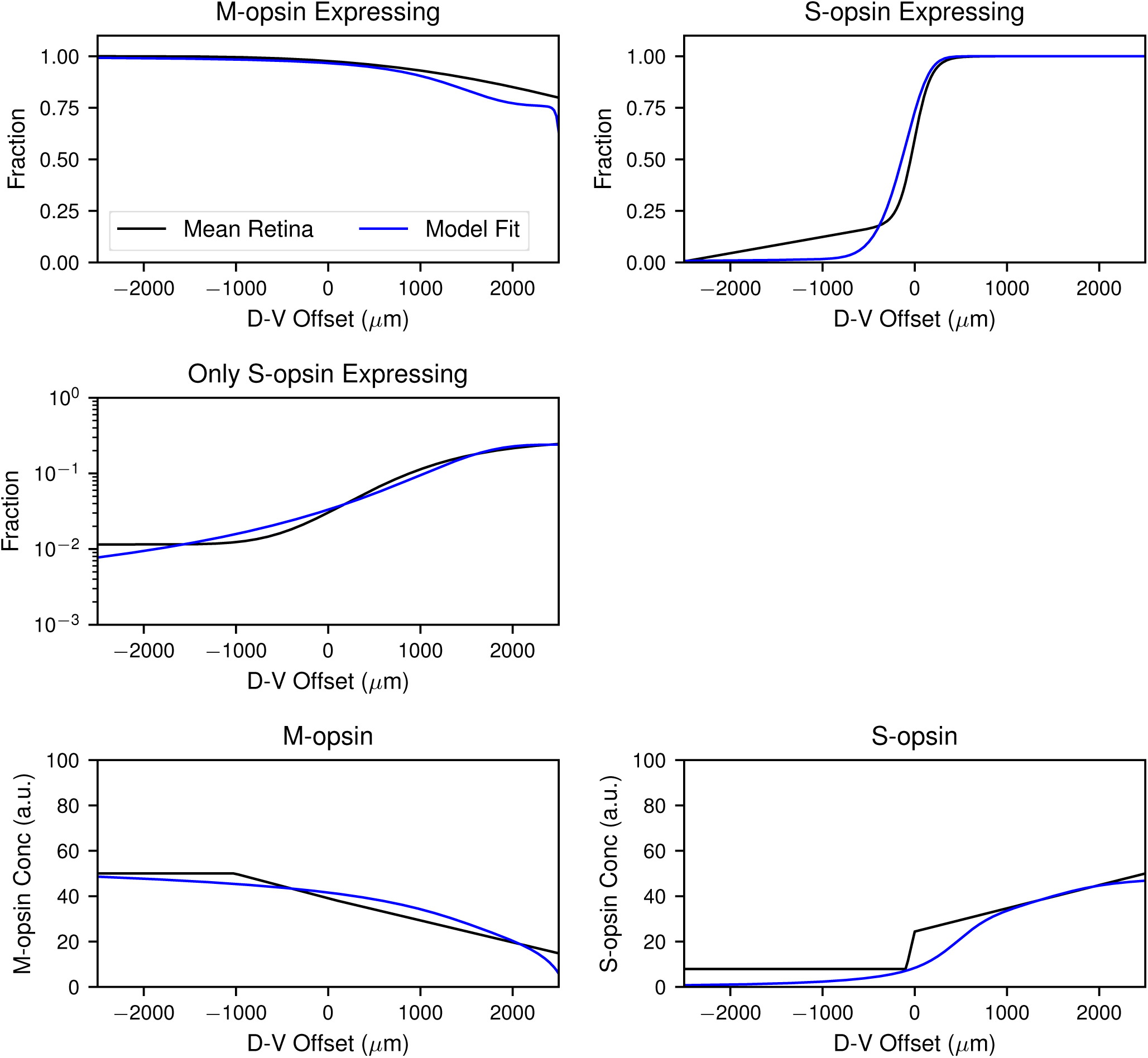
Best fit parameterization. Comparison of the best fit model parameterization (blue) with the hypothetical mean retina (black). The top row shows a comparison of the fraction of cells expressing M- and S-opsin, respectively. The middle row shows the fraction of FD(S) cells. The bottom row shows the mean M- and S-opsin concentration per cell, respectively.

